# Reactivating conflicting evaluative memories during sleep reduces decision ambivalence

**DOI:** 10.64898/2026.03.11.710434

**Authors:** Danni Chen, Tao Xia, Jing Liu, Yuqi Zhang, Xibo Zuo, Haiyan Wu, Cora S.W. Lai, Xiaoqing Hu

## Abstract

Memory guides everyday evaluations and decision-making. Yet people often encounter inconsistent information about the same target, giving rise to conflicting evaluative memories and decision ambivalence. Decision ambivalence is not only aversive but also reduces confidence, increases hesitation, and leads to maladaptive choices. While sleep consolidates memories, its role in resolving these evaluative conflicts and shaping decision dynamics remains unknown. Here, we investigated how memory reactivation during sleep, a critical period for memory consolidation and transformation, would reconstruct conflicting evaluative memories and thus influence next-day decision ambivalence. In a valence reversal learning procedure, participants first encoded positive or negative cue-outcome associations (A-B) on Day 1, followed by learning A-C associations yet with opposite valences on Day 2 (i.e., Day 1 negative-to-Day 2 positive and Day 1 positive-to-Day 2 negative). During subsequent non-rapid eye movement (NREM) sleep on Day 2 night, half of the cues were re-presented to sleeping participants to reactivate the cue-associated memories. Upon waking up, participants completed post-sleep evaluation and memory tests. Our results showed that cueing reduced decision ambivalence, especially in the negative-to-positive condition, as evidenced by less curved mouse-tracking trajectories. Meanwhile, cueing promoted the integration of conflicting evaluative memories, again in the negative-to-positive condition. Critically, cueing-induced ambivalence reduction was evident only for items that were integrated after sleep. Electrophysiologically, stronger cue-elicited delta power during NREM sleep predicted next-day ambivalence reduction, while higher cue-elicited spindle probabilities were associated with better memory integration. Together, our findings suggest that memory reactivation during post-learning NREM sleep actively reorganizes conflicting memories, providing a mechanistic pathway through which offline memory reprocessing resolves waking decision ambivalence.

## Introduction

Memory guides decision-making. The choices we make are shaped by what we have learned and how we feel (Biderman et al., 2020; Shohamy & Daw, 2015). Yet in today’s “infodemic” environment (Zarocostas, 2020), people are frequently exposed to conflicting information about the same option. For example, an initially positive impression toward a job candidate may later be disconfirmed as negative, and vice versa. Holding conflicting evaluative memories can produce attitudinal ambivalence, eliciting aversive feelings and dissonance, and lead to maladaptive choices such as substance abuse and vaccination hesitancy (Béna et al., 2022; Foster et al., 2014; Kim et al., 2019; Menninga et al., 2011; Oser et al., 2010, Schneider & Schwarz, 2017; van Harreveld et al., 2009). While decision ambivalence often arises from conflicting evaluative memories, it remains unclear how offline memory reorganization can resolve conflict and shape next-day decision-making.

A plausible, hitherto untested, hypothesis is that offline sleep memory reactivation can reduce next-day decision ambivalence. During sleep, memory undergoes systematic transformation via memory reactivation and systems-level consolidation processes (Brodt et al., 2023; Paller et al., 2021). Beyond memory consolidation and stabilization, accumulating evidence suggests that sleep transforms memory by integrating related episodes and extracting schematic knowledge (Cowan et al., 2020; Liu et al., 2025). Such sleep-dependent memory reprocessing may organize conflicting, fragmented information into coherent, structured knowledge representations that would facilitate next-day decision-making (Son et al., 2024).

Employing a paradigm known as targeted memory reactivation (TMR), researchers can re-present sensory cues related to prior daytime learning to reactivate cue-specific memories during sleep (Hu et al., 2020; Rasch et al., 2007; Rudoy et al., 2009). Via manipulating covert reactivation processes during sleep, researchers can then examine how such reactivation impacts subsequent memory, emotion, and decision-making (Temudo & Albouy, 2024; Xia & Hu, 2025). Of particular relevance to the current study, cueing can reactivate multiple pieces of memory during sleep (Schechtman et al., 2021), facilitating memory competition or integration depending on pre-sleep memory performance and the relationship among to-be-reactivated memories (Antony & Schechtman, 2023; Tamminen et al., 2010; Xia et al., 2024).

Once conflicting evaluative memories are integrated to form coherent memory structures, they allow evaluative information to be pre-computed before the decision-making, thereby reducing subsequent decision ambivalence. Indeed, recent work in memory-based decision making demonstrates that forming associative memory structures can help reward value being transferred to non-rewarded items, with the strength of association predicting the magnitude of transferred value (Biderman et al., 2023; Biderman & Shohamy, 2021; Wang et al., 2019; Wimmer & Shohamy, 2012). Complementing the behavioral evidence, multivariate neural decoding analysis reveals that neural patterns corresponding to decision-relevant memories can be evident before choices are made, consistent with the idea that the brain retrieves relevant past experiences to support decision-making (Nicholas et al., 2025; Ou et al., 2025). Overall, these converging findings suggest that the decision is shaped not only by what has been learned but also by how evaluative memories are organized and retrieved during decision-making.

Linking sleep-dependent memory transformation and memory-based decision making, we hypothesized that reactivating conflicting evaluative memories during non–rapid eye movement (NREM) sleep can facilitate memory reorganization to reduce next-day decision ambivalence. To induce conflicting evaluative memories and decision ambivalence, we employed an evaluative learning and valence reversal task, in which participants first learned stimuli pairings of positive and negative emotional valence, followed by a valence reversal learning. To provide a comprehensive examination of valence reversal effects, we included both negative-to-positive and positive-to-negative valence reversal conditions. In the negative-to-positive condition, participants learned pseudoword–negative image pairings on Day 1 evening, followed by pseudoword-positive image pairings on Day 2 evening, with the pseudowords being used as the conditioned stimuli. In the positive-to-negative condition, participants learned positive information in pseudoword–positive image pairings on Day 1 evening, followed by learning pseudoword-negative image pairings on Day 2 evening. For Day 2 NREM sleep TMR, we re-presented half of the pseudoword cues from both valence reversal conditions (i.e., cued), whereas the remaining cues were not replayed (i.e., uncued). Because each pseudoword was associated with conflicting evaluative information across two days, reactivating the conflicting evaluative memories during Day 2 NREM sleep may promote either integration or competition, leading to less or more ambivalence (Antony & Schechtman, 2023; Hennies et al., 2016; Xia et al., 2024). To quantify real-time decision ambivalence, we examined mouse-tracking trajectories when participants used the computer mouse to decide whether the pseudoword cue would lead to positive or negative consequences (Béna et al., 2022; Melnikoff et al., 2021; Schneider & Schwarz, 2017; Xu et al., 2025).

To further delineate the electrophysiological mechanisms that support sleep-dependent memory reactivation and next-day decision-making, we investigated cue-elicited sleep neural activities implicating memory reactivation. Mounting research demonstrates that cue-elicited delta Electroencephalogram (EEG) power is associated with memory reactivation and can predict post-sleep evaluation and decisions (Ai et al., 2018; Chen et al., 2024; Creery et al., 2015). Of particular relevance, cueing multiple memories increased cue-elicited delta power than cueing a single memory (Schechtman et al., 2021), suggesting that delta power may track the amount of memories being reactivated via cueing. In addition to delta power, the 12-16 Hz spindles are canonical sleep neural oscillations that mediate memory reactivation: spindle activities track item- or category-specific neural representations predicting post-sleep memory performance (Cairney et al., 2018; Liu et al., 2023).

More importantly, spindles have been intensively studied in the context of memory integration (Cowan et al., 2020; Hennies et al., 2016; Tamminen et al., 2010), and has been related to cueing-induced changes in memory and evaluation (Chen et al., 2024; Xia, Antony, et al., 2023). Building on this rich literature, we focused on how cue-elicited delta and spindle activity would support the cueing-induced changes in decision ambivalence and in evaluative memories.

## Results

Following exclusion (i.e., unsuccessful evaluative learning or failures in complying with task instructions), 42 participants (34 females; Age: *Mean* = 23.50, *S.D.* = 2.47). were included in the EEG analyses, and 36 in the behavioral analyses (28 females; Age: *Mean* = 23.61, *S.D.* = 2.51; for details, see Methods). Included participants completed the following sessions: Day 1 evaluative learning, Day 2 counter-evaluative learning, sleep-based targeted memory reactivation (TMR), and Day 3 post-TMR evaluation and memory tests (**Figure 1**).

**Figure 1.**
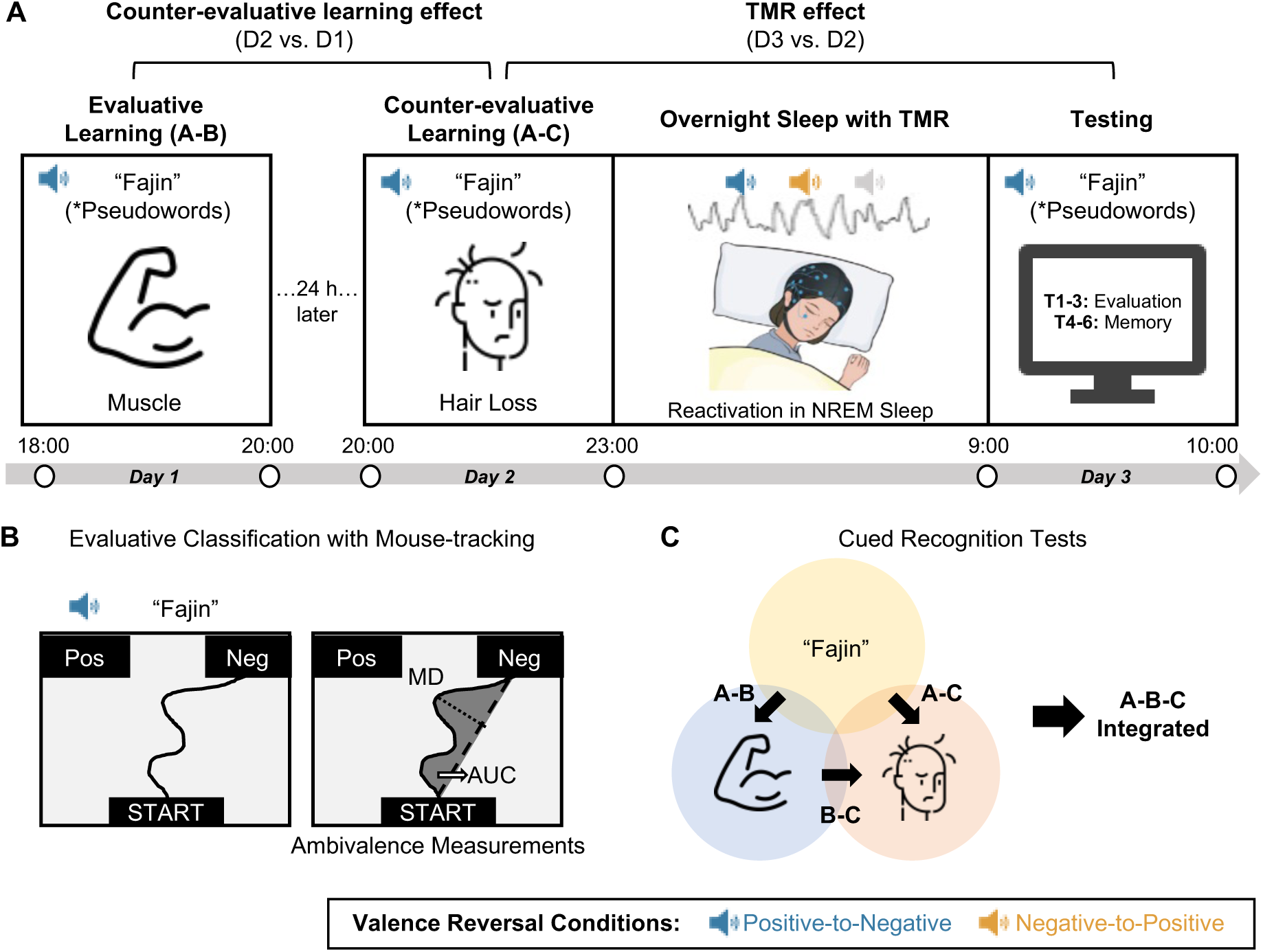
Experimental procedure and tasks. **(A) Overview.** The experiment comprised two laboratory visits. On Day 1, participants learned A (cue)-B (first health outcome) associations. On Day 2, they learned A-C (second health outcome) associations, with C having the opposite valence of B, i.e., counter-evaluative learning, constructing two valence reversal conditions: Day 1 positive-to-Day 2 negative and Day 1 negative-to-Day 2 positive. On Day 2 night, half of the cues (“A”) were acoustically presented during NREM sleep. The following Day 3 morning, participants completed a battery of evaluation and memory tests. **(B) Evaluative Classification Task.** Participants used a computer mouse to indicate whether each cue (A) led to either positive or negative health outcomes. Mouse trajectories were analyzed as an index of decision ambivalence, with larger deviations and more curved trajectories indicating higher ambivalence. **(C) Memory tests and integration criterion.** Memory was assessed with three separate cued-recognition tasks: B-C, A-B, and A-C cued recognition. Items were classified as A-B-C integrated if participants responded correctly on all three recognition tasks. The order of the A-B and A-C tests was counterbalanced across participants.

### Effective evaluative learning and counter-evaluative learning

We first verified that the evaluative learning and counter-evaluative learning procedures successfully established evaluative memories and updated evaluative decisions accordingly. On Day 1, participants successfully learned the cue-target evaluative memories, showing high recognition accuracies of A-B pairings (mean ± S.D., 0.94 ± 0.09) and highly significant valence effects in all evaluation tests, including speeded judgment, explicit rating, and evaluative classification task (*p*s < .001). No other valence reversal or TMR effects were significant, suggesting both memory and evaluations were comparable between cued and uncued conditions before TMR (*p*s > .154; **Data S1**).

On Day 2, participants successfully learned the counter-evaluative pairings, as evidenced by high A-C recognition accuracies (mean ± S.D., 0.84 ± 0.18). Moreover, participants showed higher A-C recognition memories in the negative-to-positive than the positive-to-negative valence reversal condition (*F* (1, 35) = 6.44, *p* = .016, η^2^_*G*_ = .029). Critically, counter-evaluative learning effectively updated evaluative choices and classification per the reversal conditions (*p*s < .001). No other effects were significant (*p*s > .176; **Data S2**).

### Sleep reactivation reduced next-day decision ambivalence

We next examined whether reactivating conflicting evaluative memories during NREM sleep via TMR reduces decision ambivalence. During NREM sleep on the night of Day 2, we re-presented half of the cues to reactivate cue-specific memories. On both Day 2 evening (pre-sleep/TMR) and Day 3 morning (post-sleep/TMR), participants completed evaluative classification tasks in which they moved a mouse to evaluate the cues (i.e., “positive” or “negative”). Decision ambivalence was quantified by the mouse-tracking trajectories in terms of Area Under the Curve (AUC), Maximum Deviation (MD), and Average Deviation (AD). We then summed these measures to obtain a comprehensive index of decision ambivalence, with larger values indicating higher decision ambivalence (Figure 1B, see Methods for details; results for individual measurements are provided in **Data S3**).

We fitted item-level Bayesian linear mixed models (BLMMs) with TMR (cued vs. uncued), valence reversal (negative-to-positive vs. positive-to-negative), and time (Day 2 pre-TMR vs. Day 3 post-TMR) as fixed effects. The model showed a credible TMR by time interaction (Median_diff_ = 0.18, 95% HDI [0.03, 0.33]). Post-hoc analyses indicated that cueing reduced ambivalence from Day 2 pre-TMR to Day 3 post-TMR (pre-vs. post-TMR, Median_diff_ = 0.09, 95% HDI [0.00, 0.18]), while uncued items showed no credible changes (Median_diff_ = -0.03, 95% HDI [-0.12, 0.07]; **Figure 2A**).

**Figure 2.**
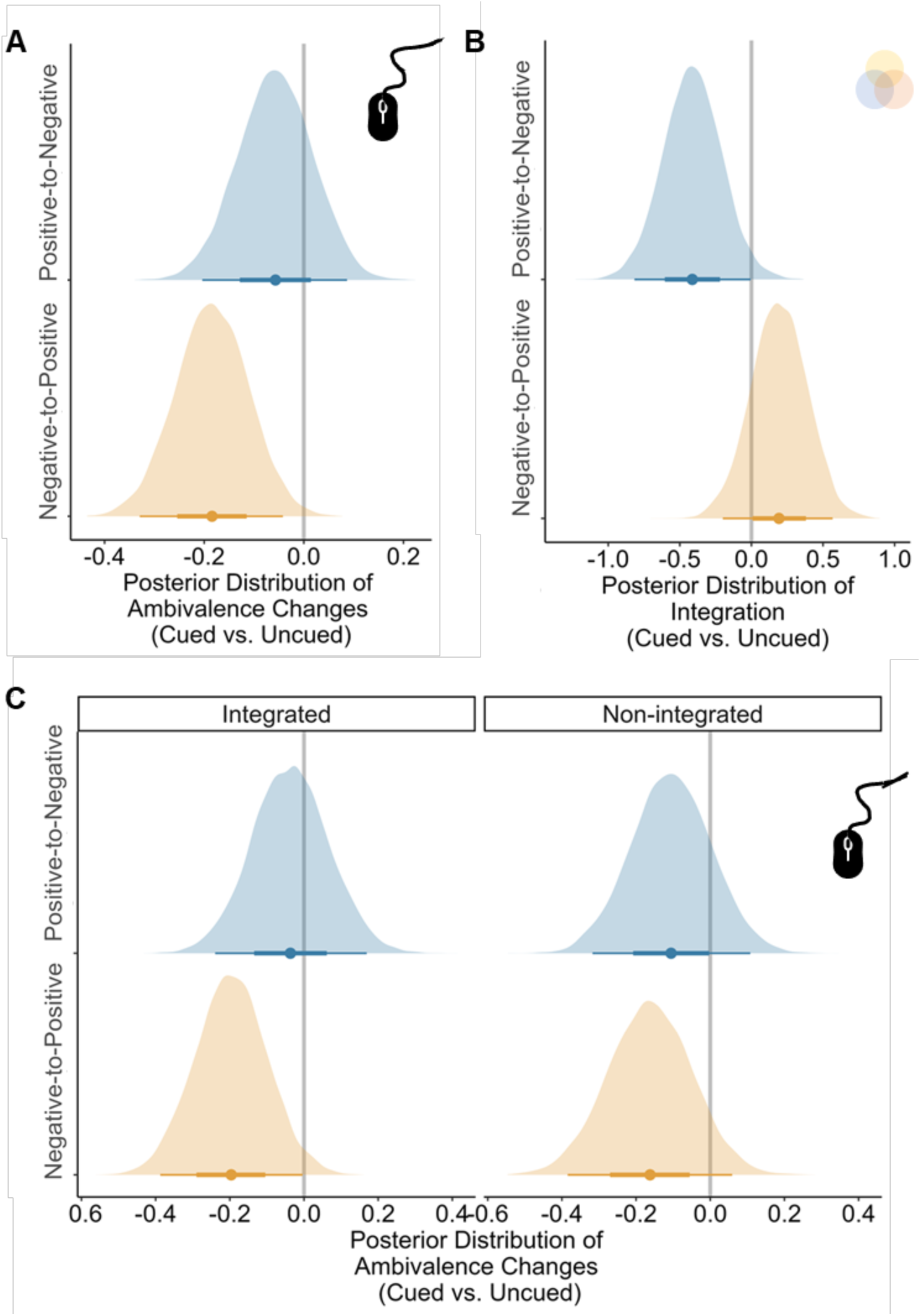
Bayesian posterior estimates of TMR effects on decision ambivalence and memory integration. **(A)** Posterior distributions of the cued minus uncued difference in ambivalence change (Day 3 post-TMR minus Day 2 post-counter-evaluative learning), shown separately for the two valence-reversal conditions (negative-to-positive, orange; positive-to-negative, blue). **(B)** Posterior distributions of the cued-uncued difference in post-TMR A-B-C triplet integration, plotted separately for each valence-reversal condition. **(C)** Posterior distributions of the cued minus uncued difference in ambivalence change (Day 3 post-TMR minus Day 2 post-counter-evaluative learning), shown separately for items classified as post-TMR integrated (left) versus non-integrated (right), and for each valence-reversal condition. In each panel, the shaded density shows the posterior distribution, the dot indicates the median, and the horizontal segment indicates the 95% highest-density interval (HDI). The vertical reference line at 0 denotes no cued minus uncued difference; effects were considered reliable when the 95% HDI did not include 0.

Focal analyses examining each valence reversal condition revealed that the cueing credibly reduced ambivalence from Day 2 pre-TMR to Day 3 post-TMR in the negative-to-positive valence reversal condition (cued vs. uncued, Median_diff_ = -0.18, 95% HDI [-0.33, -0.04]). On the contrary, no cued vs. uncued effects were found in the positive-to-negative valence reversal condition, Median_diff_ = -0.06, 95% HDI [-0.20, 0.09]). Overall, these findings suggest that reactivating conflicting evaluative memories during sleep reduced decision ambivalence, particularly in the negative-to-positive valence reversal condition.

We also found that counter-evaluative learning continuously updated explicit evaluative ratings from Day 1 post-evaluative learning to Day 3 post-TMR tests: the negative-to-positive reversal condition showed higher rating changes than the rating changes in the positive-to-negative condition (*p* < .001). However, neither the TMR effect nor its interaction was significant (all *p*s > .231; **Data S4**).

### Sleep reactivation impacts the integration of conflicting evaluative memories

To probe the memory mechanisms underlying the TMR-related change in decision ambivalence, we next tested how TMR influenced evaluative recognition memory for (i) individual A-B and A-C pairs and (ii) integrative A-B-C memory structure. Building on prior relevant work (Cox et al., 2021; van Kesteren et al., 2018), we derived an integration memory score: each A-B-C triplet was classified as integrated if the participant responded correctly to all three individual tests (A-B, A-C, and B-C pairs); otherwise, it was classified as non-integrated (**Figure 1C**).

While TMR did not affect A-B or A-C memories (all 95% HDI includes 0; **Data S5**), we found TMR effects on A-B-C triplet memory integration depending on valence reversal (**Figure 2B**). An item-level BLMM taking valence reversal and TMR as fixed factors revealed a credible TMR by valence reversal interaction on integration (Median_diff_ = 0.61, 95% HDI [0.07, 1.17]). Post-hoc analyses showed that cuing negative-to-positive items enhanced integration more than the cueing positive-to-negative items (Median_diff_ = 0.47, 95% HDI [0.08, 0.86]), whereas this valence reversal difference was not credible among uncued items (Median_diff_ = -0.14, 95% HDI [-0.56, 0.28]). Furthermore, within the positive-to-negative condition, cueing reduced integration (cued vs. uncued, Median_diff_ = -0.41, 95% HDI [-0.81, 0.00]). The results remained the same after controlling for the within-pair absolute valence rating differences (**Data S6**).

#### Pre-sleep memory performance matters for TMR-induced integration

To better understand how pre-sleep memory performances influence TMR-induced memory changes (Cairney et al., 2016; Creery et al., 2015; Schechtman et al., 2023), we quantified pre-sleep/TMR memory performance on integration. Given no B-C memory test was performed before TMR, we scored items as pre-TMR integrated if participants responded correctly to both Day 1 A-B and Day 2 A-C recognition tests; otherwise, they were identified as pre-sleep non-integrated.

We conducted a BLMM with valence reversal and TMR as fixed factors on pre-sleep memory integration. Similarly, the negative-to-positive condition had higher integration than the positive-to-negative condition (Median_diff_ = -0.54, 95% HDI [-1.03, -0.04]). This effect remained the same after controlling for the absolute valence rating difference within each pair (Median_diff_ = -0.52, 95% HDI [-1.01, -0.01]). In addition, no credible TMR effect nor their interaction was found, suggesting the pre-TMR performance did not credibly differ between to-be-cued and to-be-uncued items (i.e., successful randomization, all corresponding 95% HDI included 0).

To further understand the relationship between pre- and post-TMR integration, we divided items into pre-sleep integrated versus pre-sleep non-integrated, and fitted two BLMMs with valence reversal and TMR as fixed factors on post-TMR A-B-C triplet integration. The critical valence reversal by TMR interaction emerged only among the pre-integrated items (Median_diff_ = 0.75, 95% HDI [0.12, 1.40]). Post-hoc analyses again showed divergent TMR effects for the two valence reversal conditions: TMR cueing numerically enhanced the post-TMR integration for the negative-to-positive valence reversal condition (cued vs. uncued, Median_diff_ = 0.29, 95% HDI [-0.18, 0.75]), but numerically reduced the post-TMR integration for the positive-to-negative condition (cued vs. uncued, Median_diff_ = -0.45, 95% HDI [-0.95, 0.01]). In contrast, no credible interaction effect was observed for non-pre-integrated items (Median_diff_ = -0.70, 95% HDI [-3.06, 1.11]). These results further suggest that TMR effects on memory integration (A-B-C) emerged primarily for items that were already partially integrated before sleep (A-B and A-C).

### Ambivalence reduction among post-TMR integrated items

Reactivating counter-evaluative learning memories during sleep reduced the next-day decision ambivalence and preserved memory integration, particularly in the negative-to-positive valence reversal condition. We next asked whether ambivalence reduction was specifically associated with post-TMR memory integration. To test this, we fitted a BLMM predicting ambivalence with TMR (cued vs. uncued), valence reversal (negative-to-positive vs. positive-to-negative), session (pre-vs. post-TMR), and post-TMR integration status (integrated vs. non-integrated) as fixed factors.

Although there were no credible interactions (Median_diff_ = 0.10, 95% HDI [-0.31, - 0.52]), focal analyses showed that cueing-induced ambivalence reduction only emerged among post-TMR integrated items in the negative-to-positive condition (cued vs. uncued, Median_diff_ = -0.20, 95% HDI [-0.39, -0.01]; **Figure 2C** left). In contrast, no reliable pre-vs. post-TMR changes were observed in any other conditions (cued vs. uncued, -0.16 <= Median_diff_ <= -0.04, all 95% HDIs included 0; **Figure 2C** right). Together, these results suggest that the cueing-induced ambivalence reduction was particularly evident among integrated memories, particularly in the negative-to-positive reversal condition.

### Cue-elicited delta and spindle power predicted ambivalence and integration

Cue-elicited delta and spindle activities support memory and evaluation (Ai et al., 2018; Chen et al., 2024; Xia, Antony, et al., 2023). We next examined how cue-elicited delta/spindles impact decision ambivalence and memory integration.

#### Cueing elicited significant delta and spindle changes

We first characterized cue-elicited EEG power changes during sleep. Relative to the [-1000 to -200 ms] pre-cue baseline, re-playing memory cues significantly enhanced the broadband 1-30 Hz power during an early cluster (−60 to 3272 ms relative to the cue onset, *p*_cluster_ = .001, cluster-based permutation test corrected; see Methods for details), but reduced the 10.5 to 17.0 Hz power in a later cluster (2200 to 4108 ms, *p*_cluster_ = .024; **Figure 3A**). Similarly, the control, memory-irrelevant cues enhanced the 1-30 Hz EEG power in the early cluster [40 to 2868 ms] (*p*_cluster_ = .001) but reduced the 10.5 to 16.5 Hz power in the later cluster [2188 to 3652 ms], *p*_cluster_ = .035 (**Figure 3B**).

**Figure 3.**
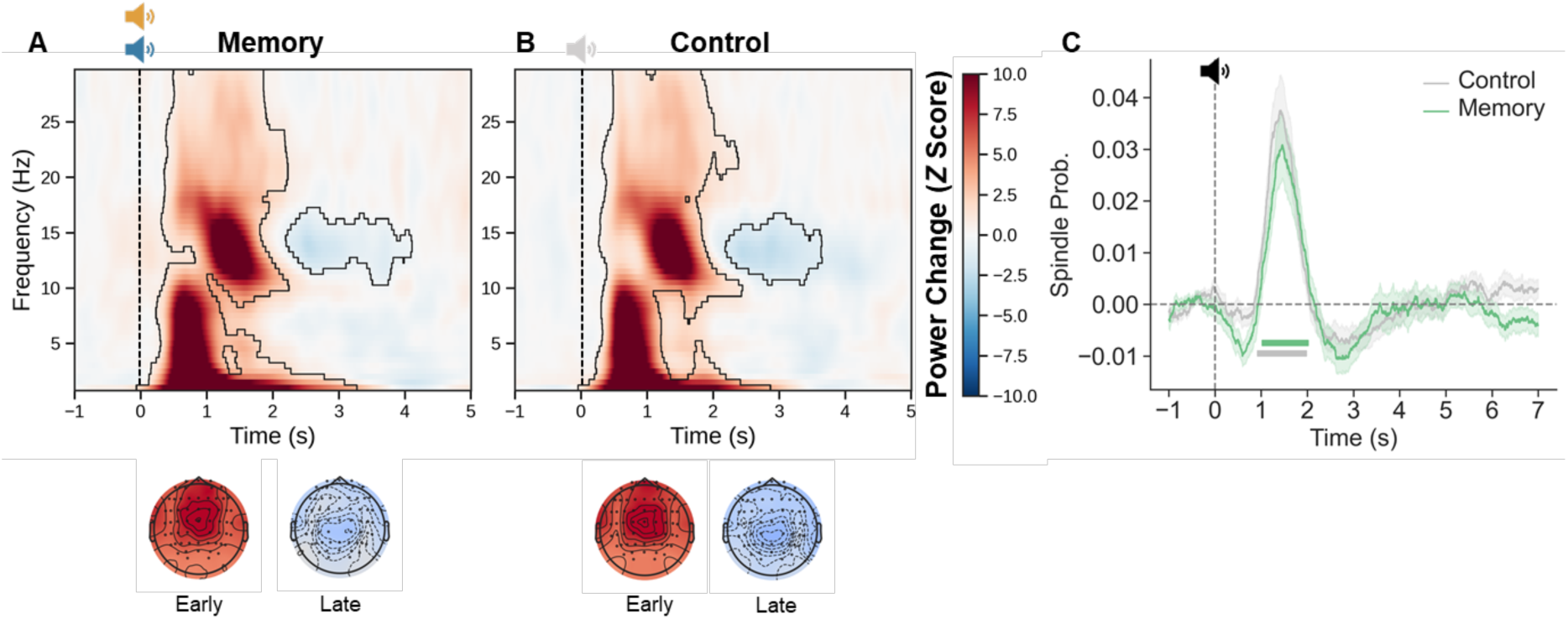
TMR Cued-elicited EEG Activities. Both **(A)** memory and **(B)** control cues elicited spectral power changes. **(C)** Both memory and control cues elicited significant spindle probability changes. The colored line below indicates the significant cluster comparing cue-elicited spindle probability and baseline. The black line below indicates the significant cluster comparing all trials with the baseline.

Regarding spindle activities, compared to the -1000 to 0 ms pre-cue baseline, both memory cues and control cues significantly increased the spindle probability (memory cues, 1024 - 2020 ms, *p*_cluster_ = .014; control cues, 920 - 1980 ms, *p*_cluster_ = .001; **Figure 3C**), with no significant differences between memory and control cues (cluster-based permutation test, *p*_cluster_ = .450). Overall, while cueing reliably elicited significant EEG activity, no differences were found between memory or control cues nor between different valence reversal conditions.

### Cue-elicited delta power tracks ambivalence reduction

Upon confirming that memory cues elicited significant delta and spindle activity changes, we next investigated how cue-elicited EEG activities predicted ambivalence reduction from pre- to post-TMR using BLMM, with cue-elicited delta power/spindle probability and valence reversals as fixed factors and the ambivalence changes from pre-TMR to post-TMR as the dependent factor.

We found a credible interaction effect between delta power and valence reversal on ambivalence reduction (Median_diff_ = 0.02, 95% HDI [0.00, 0.04]). Post-hoc analysis showed that the relationship between delta power increase and ambivalence reduction was credibly more negative in the negative-to-positive condition (Median_negative-to-positive_ = -0.01, 95% HDI [-0.02, 0.00]) than that of the positive-to-negative condition (Median_positive-to-negative_ = 0.01, 95% HDI [-0.01, 0.03]; **Figure 4A**). The results indicated a stronger delta increase - ambivalence reduction correlation in the negative-to-positive condition than in the positive-to-negative valence reversal condition. Thus, cue-elicited delta activity tracked the extent of ambivalence reduction selectively in the negative-to-positive reversal condition.

**Figure 4.**
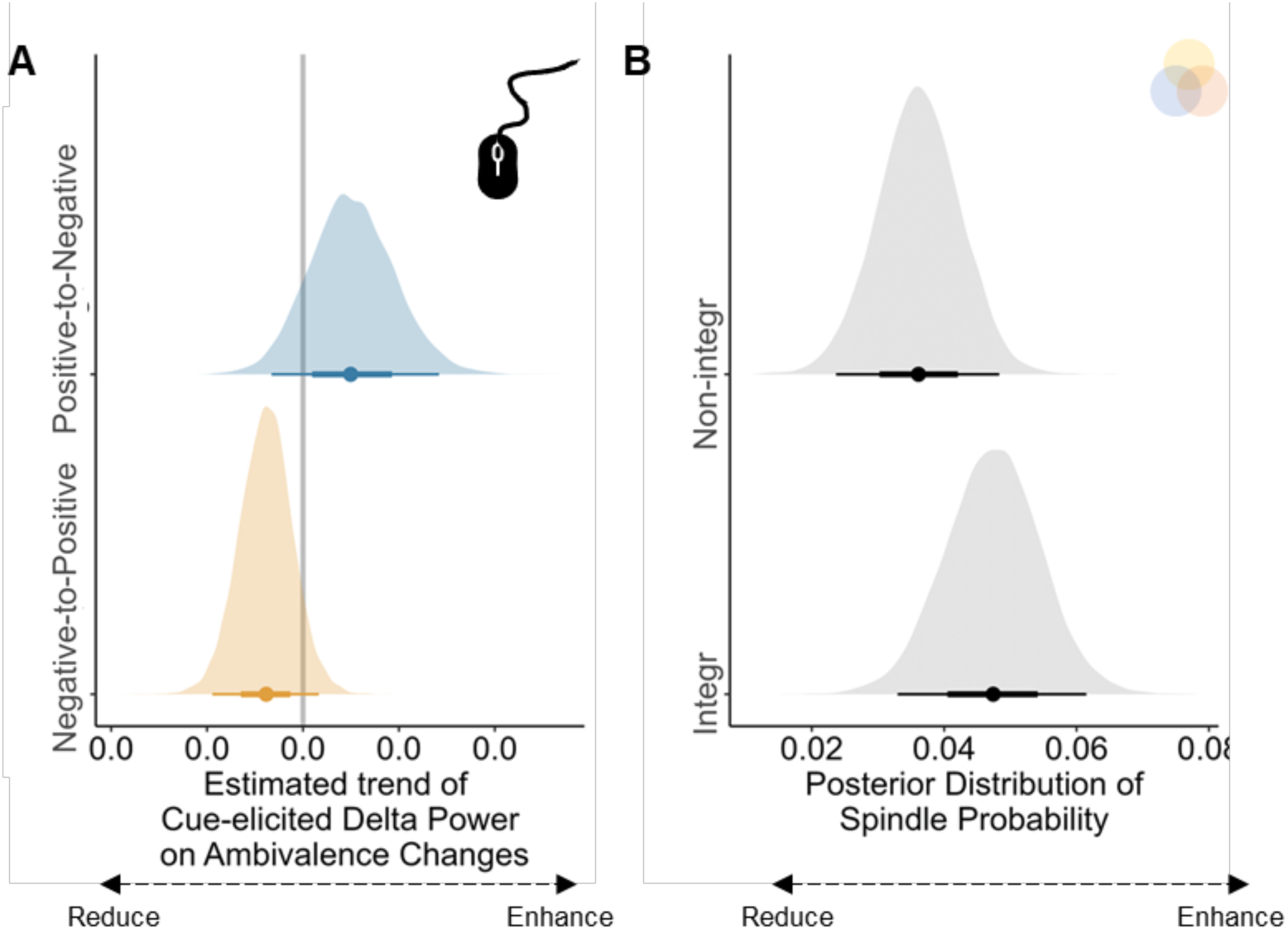
EEG-Behavior relationships. **(A)** Posterior distributions of the regression slope relating cue-elicited delta power to changes in decision ambivalence (Post-TMR minus Pre-TMR), shown separately for the Positive-to-Negative (blue) and Negative-to-Positive (orange) reversal conditions. **(B)** Posterior distributions of cue-elicited spindle probability, shown separately for items classified as integrated versus non-integrated. In both panels, shaded densities indicate posterior distributions; dots mark posterior medians; horizontal bars indicate 95% highest-density intervals (HDIs). Vertical reference lines at 0 denote no effect.

By contrast, spindle probability did not predict ambivalence reduction (−2.34<Median_diff_ < 0.95, all 95% HDIs include 0).

### Cue-elicited spindle probability tracks memory integration

We next adopted BLMM with cue-elicited spindle power/probabilities as a dependent variable, with post-TMR integration and valence reversal as fixed factors. Results showed that cue-elicited spindle probability (1024 - 2020 ms) was credibly higher for integrated than for non-integrated triplets (Median_diff_ = 0.01, 95% HDI [0.00, 0.02]; **Figure 4B**). However, there was no credible difference in cue-elicited delta power between integrated vs. non-integrated items (Median_diff_ = -0.25, 95% HDI [-1.81, 2.33]).

Overall, these item-level EEG-behavioral correlational analyses suggest that sleep-dependent memory reactivation processes, as indicated by cue-elicited delta power and spindle probabilities, contributed to cueing-induced ambivalence reduction and memory integration.

## Discussion

In today’s “infodemic” (Zarocostas, 2020), people frequently encounter inconsistent information that leads to conflicting evaluations and thus decision ambivalence. Decision ambivalence undermines confidence, delays responses, and causes maladaptive outcomes, including indecision, procrastination, and risky choices, among others (Schneider & Schwarz, 2017). Here, we provided new evidence that reactivating conflicting evaluative memories via targeted memory reactivation (TMR) during non-rapid eye-movement (NREM) sleep reduced next-day decision ambivalence, as evidenced by real-time mouse tracking trajectories. During TMR, cue-elicited delta activity tracked individual differences in ambivalence reduction, with cue-elicited spindle activity supporting the integration of conflicting memories.

Together, these findings connect sleep-mediated memory reprocessing to the real-time resolution of decision ambivalence, shedding light on how sleep memory reactivation and reorganization can support adaptive decision-making.

A key contribution of our study is that we document how sleep memory reactivation would benefit next-day decision making via the lens of memory reorganization and dynamic decision ambivalence (Béna et al., 2022). We found that TMR reduced decision ambivalence during evaluative classification, as evidenced by less curved mouse trajectories. Instead of resolving ambivalence via conscious efforts and intentional control (Béna et al., 2022; Freeman & Ambady, 2010; Lim et al., 2018), we provided an alternative route: sleep-mediated reactivation and reorganization of conflicting evaluative memories may also reduce decision ambivalence. Our results are in line with prior work showing that sleep-based reactivation can facilitate memory reorganization and benefit subsequent behavior (del Río et al., 2026; Durrant et al., 2011; Ellenbogen et al., 2007; Lewis & Durrant, 2011; Siefert et al., 2024). Supporting the role of memory reactivation in resolving decision ambivalence, stronger cue-elicited delta EEG power (1 - 4 Hz) predicted reduced magnitude of ambivalence in the negative-to-positive condition. This aligns with prior TMR work linking delta activity to evaluation updating (Ai et al., 2018; Chen et al., 2024; Xia et al., 2024) and with broader evidence that delta activity supports memory consolidation (Cho et al., 2025; Creery et al., 2015; Rihm et al., 2014; Yuksel et al., 2025). Stronger delta activity may reflect more effective cue-triggered reactivation, potentially involving the reinstatement of multiple interrelated memories (Schechtman et al., 2021), thereby helping resolve competition between conflicting evaluative tendencies and reducing next-day decision ambivalence.

A plausible mechanism through which sleep-dependent reactivation reduces ambivalence is by reorganizing conflicting evaluative memories. Specifically, re-playing the overlapping memory cues (“A”) may facilitate the integration of two episodes (A-B and A-C) into a relational A-B-C representation. Memory integration may then allow valence information to be “pre-computed” and retrieved more efficiently during later evaluation (Nicholas et al., 2025; Ou et al., 2025; Shohamy & Daw, 2015). Consistent with this account, although TMR did not improve accuracy on any single cued-recognition test, it selectively modulated A-B-C integration depending on valence reversal and pre-sleep integration status. Specifically, TMR preserved integration in the negative-to-positive valence reversal condition - the same condition in which TMR reduced decision ambivalence. Critically, the cueing-induced reduction in ambivalence was most pronounced for items that showed successful post-sleep A-B-C integration. Supporting this memory integration mechanism, our spindle results provide a plausible neurophysiological underpinning: higher cue-elicited spindle probability was associated with greater post-TMR A-B-C integration. Our finding thus extends prior work that implicates spindles in sleep-dependent memory integration to memory-based decision-making (Cowan et al., 2020; Fernandez & Lüthi, 2020; Tamminen et al., 2010). Specifically, spindles constitute a key memory reprocessing window during which newly encoded memories are reactivated and reorganized (Antony et al., 2019). In TMR, cue-elicited spindle-related activity tracked categorical and even item-specific memory representations that predicted memory consolidation(Abdellahi et al., 2026; Cairney et al., 2018; Liu et al., 2023). In our study, cue-locked spindles may index moments when multiple competing evaluative memories were reactivated and integrated into a coherent relational structure, thereby reducing competition between conflicting evaluations and ultimately lowering next-day decision ambivalence.

Critically, we observed an asymmetrical, yet consistent effect across different valence reversal conditions in both memory integration and decision ambivalence. Cueing preserved integration for negative-to-positive items, yet reduced integration in the positive-to-negative condition, whereas. This pattern is consistent with the theoretical accounts that TMR effects on interlinked memories depend on the pre-sleep relationship between the memories: cueing promotes integration when overlapping memories were encoded in a harmonious context, but can disrupt memory when they were encoded in an antagonistic context (Antony & Schechtman, 2023). Our cover story emphasized that each hypothetical pharmaceutical product (cue) could produce both desirable outcomes (one of the targets) and aversive side effects (another target). Specifically, the negative-to-positive sequence may have rendered the second (positive) information more desirable and more expected given the context (i.e., pharmaceutical product shall yield desirable health outcomes), leading to better memories and integration than the positive-to-negative condition. In line with this argument, we found that the pre-sleep integration was already stronger for the negative-to-positive than the positive-to-negative valence reversal condition, suggesting a more optimal encoding before sleep in the negative-to-positive valence reversal order. The preferential processing of positive evaluation in the negative-to-positive condition is consistent with the established literature in the optimistic belief updating, wherein people preferentially encode and consolidate desirable than undesirable feedback in belief updating (Eil & Rao, 2011; Sharot & Garrett, 2016; Yao et al., 2021). Future research could directly manipulate the desirability of Day-2 information while keeping the valence consistent to investigate how desirability influences the reversal learning effect.

Despite effects on ambivalence dynamics and integration indices, we did not observe robust TMR effects on recognition accuracy in any single recognition memory measure. Several factors may account for these null findings. First, recognition performance was already at ceiling before sleep, potentially preventing further TMR benefits in recognition memory. Second, TMR outcomes are sensitive to the memory measure employed (Hu et al., 2020): as in prior work, cued-recognition tasks do not consistently reveal TMR-related benefits even there is an observable memory representation during sleep (Ashton et al., 2018; Cairney et al., 2016; Liu et al., 2023). Future studies may therefore benefit from more sensitive assays, such as free recall and verbal reports, and more fine-grained memory scoring including details and gist (Küpper et al., 2014; Xia et al., 2024). Finally, memory reorganization can unfold over extended timescales (Dudai et al., 2015), such that TMR-related benefits (or costs) become apparent only after longer delays (Abdellahi et al., 2026; Cairney et al., 2018; Oudiette & Paller, 2013; Rakowska et al., 2021). Incorporating delayed tests would help determine whether TMR effects on single-item memory emerge over time.

Limitations should be acknowledged. First, we focused on exogenous reactivation during NREM sleep (Hu et al., 2020). However, previous research and theoretical models also suggest that REM sleep contributes to integration and schematic transformation (Cai et al., 2009; Lewis et al., 2018; Liu et al., 2025; Pereira et al., 2023; Tamminen et al., 2017). Future studies should directly compare NREM versus REM cueing to determine the impact of different sleep stages on memory integration and decision-making. Second, although we identified sleep neural activities that support decision-making and memory, the precise mechanisms remain further investigation – was it first or second targets, or an integrated representation that was reactivated? Future research shall employ multivariate representational similarity analysis or decoding approaches with functional localizer to identify the fined-grained neural representations of reactivation and integration during sleep (Abdellahi et al., 2023; Liu et al., 2023; Schechtman et al., 2023).

To conclude, our study provides new evidence that during NREM sleep, memory reactivation can reduce next-day decision ambivalence and reorganize conflicting evaluative memories. These behavioural benefits were accompanied by cue-elicited delta and spindle activities, highlighting a role for sleep-mediated memory reactivation in supporting memory-based decision-making. Broadly, our findings contribute to the theoretical framework of memory-based decision making, suggesting that sleep can support adaptive decision-making via reorganizing and integrating mixed or rapidly changing information.

## Methods

### Participants

We recruited 58 participants from a local university (43 Females; Age, *Mean* = 23.29 years old, *S.D.* = 2.58). Participants were excluded from subsequent behavioral and EEG analysis if the memory recognition accuracy after the first day of evaluative learning was lower than 80% (*n* = 8), the cues were played fewer than three blocks (*n* = 5), or they dropped out of the experiments (*n* = 3). Six additional participants were excluded from behavioral analyses because they did not follow the experiment instructions, for example, choosing the same options across all trials. As a result, 42 participants were retained in the EEG analyses (34 females; Age: *Mean* = 23.50, *S.D.* = 2.47), while 36 were retained in the behavioral analyses (28 Females; Age, *Mean* = 23.61, *S.D.* = 2.51). All the participants were native Mandarin speakers, right-handed, not color-blind, had normal or corrected-to-normal vision, and had no reported history of neurological, psychiatric, or sleep disorders. In addition, all participants had regular sleep-wake cycles and reported good sleep quality. This research was approved by the Human Research Ethics Committee of the University of Hong Kong (HREC No. EA1904004). All participants provided written informed consent before participating in the experiment and were compensated for their participation.

### Stimuli

For materials used in the evaluative learning and counter-evaluative learning task, we used pseudowords as the names of hypothetical pharmaceutical products (i.e., cues) and images depicting either positive or negative health outcomes (i.e., targets). Each cue would be associated with two images depicting health outcomes of opposite valence and of distinct themes (i.e., hair loss vs. bright eyes).

Sixty-four two-character pseudowords were generated by randomly pairing two neutral characters from the Chinese Affective Words System (Luo & Wang, 2004; Xia et al., 2024; Xia, Yao, et al., 2023). The spoken words, which were used as the auditory memory reminders in later targeted memory reactivation (TMR) cueing, were generated in Mandarin using the Microsoft Azure Text-to-Speech function (language = “zh-CN,” duration: *Mean* ± *S.D.*, 1166.21 ± 31.54 ms). Participants rated these spoken words highly audible (percentage of choosing “audible”: *Mean* ± *S.D.*, 98.13% ± 3.60%). For each participant, we randomly selected 48 pseudowords as memory cues to be used in the evaluative learning task, while the remaining 16 pseudowords were not paired with any images and served as the non-memory control cues during the TMR.

A total of 80 health-related emotional images were selected from publicly accessible evaluative learning databases (Heycke & Gawronski, 2020), the International Affective Picture System (IAPS; Lang et al., 2005), and Google Images, with half being positive and the other half being negative. From this image set, 32 pairs were created, each consisting of one positive and one negative image, ensuring no direct semantic connection between the images to avoid potential confounding factors. For instance, a negative image depicting hair loss would not be paired with a positive image showcasing attractive hair. These pairs generated 32 unique combinations, which were randomly assigned to four experimental conditions: negative-to-positive cued, negative-to-positive uncued, positive-to-negative cued, and positive-to-negative uncued. The remaining 16 images were not paired with counter-valenced images, serving as filler stimuli (see below for details). An independent sample of participants (N = 34) rated the images on perceived healthiness and willingness to buy on a 1-9 scale. Results showed that negative images were significantly lower in perceived healthiness and willingness to buy (*p*s < .001), confirming the valence manipulation. Additionally, within-pair valence differences in healthiness and willingness to buy did not vary by TMR assignment, valence-reversal order, or their interaction, indicating that the valence of the image pairs was well matched across conditions (*p*s < .449).

### Design and Procedures

#### Overview of the Procedures

We employed a 2 (valence reversals: negative-to-positive and positive-to-negative) by 2 (TMR: cued vs. uncued) within-subject design. Participants visited the lab twice in two consecutive days (**Figure 1A**). On Day 1 evening (initial evaluative learning), participants learned the 48 pairs of pseudowords-health outcomes (either negative or positive, A-B associations). On Day 2 evening (counter-evaluative learning), participants learned 36 of the 48 pseudowords to be paired with health outcomes with opposite valence to the Day 1 learning (A-C associations), forming two valence reversal conditions: negative-to-positive and positive-to-negative. The remaining 16 pseudowords were also presented but were paired with a meaningless mosaic image, thus serving as fillers. During Day 2 non-rapid eye-movement (NREM) sleep following the counter-evaluative learning, half of the pseudowords in each valence reversal condition were randomly selected and replayed, giving rise to the cued and uncued conditions.

On Day 1 evening around 18:00, participants arrived at the laboratory, provided the written consent form and were introduced the overall procedure. Participants completed the following tasks in order: (1) Psychomotor Vigilance Task (PVT), which assessed participants’ vigilance level; (2) Pseudowords Familiarization Task, in which participants familiarized with the spoken pseudowords, to be used as the names for hypothetical pharmacological products; (3) Images Healthiness Rating Task of the images to-be-presented in the first day; (4) Evaluative Learning in which participants learned the hypothetical pharmaceutical products (i.e., pseudowords) and their associated health outcome (i.e., images, A-B associations); (5) A-B Cued

Recognition Task in which participants should choose the correct health outcome pictures, prompted by the pseudowords; (6) Speeded Choice Task in which participants made binary choices about the products (choose or not); (7) Evaluative Classification with Mouse-tracking Task, in which participants used mouse to choose the valence of the product; (8) Explicit Healthiness Rating Task, in which participants rated how healthy of the product. We employed a pre-defined learning criterion: if participants’ recognition accuracy was lower than 80% but higher than 65%, they would complete one additional round of learning and recognition task, before the speeded choice task. Participants would not proceed if they could not meet the criteria (last round of A-B Cued Recognition Task accuracy > 80%).

Qualified participants were invited to the Day 2 experiments. One Day 2 evening, participants arrived at the laboratory at around 20:00. After cleaning up and EEG setup, they completed the following tasks at around 21:30 in order: (1) PVT; (2) Baseline A-B Mental Retrieval task; (3) Images Healthiness Rating Task of the stimuli to-be-presented in the second day; (4) Counter-evaluative Learning Task in which participants learned the same products (i.e., pseudowords) yet to be paired with health outcomes with opposite valence to the Day 1 evening learning (A-C associations); (5) A-C Cued Recognition Task in which participants should choose the correct health outcome images from the counter-evaluative learning task, being prompted by the pseudowords; (6) Speeded Choice Task; (7) Evaluative Classification with Mouse-tracking. After completing these tasks, participants went to bed at around 23:30 and were given eight hours for bedtime. During participants’ sleep, experienced experimenters monitored participants’ EEG and implemented the cueing procedure.

On Day 3 morning, participants were awakened by experimenters at around 7:30 (i.e., after 8 hours bed time). After breakfast and refreshing up, participants completed the following tasks in order: (1) PVT Task; (2) Speeded Evaluative Choice Task; (3) Evaluative Classification with Mouse-tracking; (4) Explicit Healthiness Rating Task; (5) a surprise B-C Cued Recognition Task; (6) A-C and A-B Cued Recognition Task, with the order being counter-balanced across participants. These three recognition tasks were separated by a 3-minute math task for distraction, in which participants judged whether a simple mathematical equation (e.g., 4 + 5 > 7) was correct.

The details of each task are provided below. All tasks were programmed and presented by *PsychoPy* (2020.1.3; Peirce et al., 2019).

#### Psychomotor Vigilance Task (PVT)

To test whether vigilance levels might differ across different days, participants completed a 5-minute PVT each day before working on other tasks. During the PVT, a fixation was first presented on the center of the screen with a jitter duration of 2-10 s. Next, a counter starting from 0 would replace the fixation. Participants pressed the button as soon as they detected the changes. Their response times (RTs) were presented on the screen as feedback for their performance. We found no significant differences in response times across phases (*F* (1.28, 43.41) = 0.03, *p* = .904, 𝜂^2^_*G*_ < 0.001), confirming no difference in the vigilance level across experiment sessions.

#### Pseudowords Familiarization Task

Following the PVT, participants got familiarized with all 64 spoken names in the pseudowords familiarization task. Participants were told that these pseudowords represented the names of hypothetical pharmaceutical products. Each trial started with a 0.5 s fixation, followed by a pseudoword, which was presented on the center of the screen for 1 s, accompanied by its spoken name (i.e., “Fajin”) being played via an external speaker. Next, participants pressed the button to indicate whether the spoken names were clear to them. The inter-trial interval (ITI) was 1 second. The task included two blocks, each containing all 64 spoken names, being randomly presented.

#### Image Healthiness Rating Task

Participants completed the healthiness rating task twice on both experimental days. The task involved evaluating the healthiness of the central elements within the images, which could be a part of the body (e.g., legs) or a human. This healthiness rating task also served as a familiarization process for these images. Each trial started with a 0.5 s fixation, followed by the presentation of images depicting various health outcomes. Next, participants used a mouse to rate the healthiness of the image on a scale ranging from 1 (extremely bad) to 9 (extremely good). Participants rated only the images to be learned on the same days, that is, 48 images for Day 1 and 32 images for Day 2.

#### Evaluative Learning Task (A-B associative learning)

Participants would form the initial evaluation of the pharmaceutical products on the first-day evaluative learning task. We asked participants to learn 48 pairs of pseudowords (i.e., a pharmaceutical product) and their corresponding negative or positive health outcomes in the evaluative learning task (A-B associations, **Figure 1A**). Half of the health outcomes were negative, while the other half were positive. The evaluative learning task included four encoding blocks. To increase the authenticity, participants were given the following instructions at the beginning of the evaluative learning task:

*“In this task, you will be presented with names of pharmaceutical products and visual information about their effects. As you know, many pharmaceutical products have positive effects, but some products also have negative side effects. Therefore, it is important to understand pharmaceutical products and their associated outcomes for our health. For each product, you will hear their names and see how this product CAUSES health outcomes, which could be either positive or negative. Your task is to think and remember the associations, such that the pharmaceutical product CAUSES what is displayed in the outcome image.”*

In each trial of the encoding blocks, a fixation was presented for 0.5 s, followed by a blank screen with a jitter duration between 0.8 and 1.2 s. Next, a pseudoword (i.e., product names) was aurally presented, together with a corresponding health outcome image being visually presented on the screen for 3 s. Participants were required to think and memorize the pseudoword-outcome associations during this encoding phase. After that, the outcome image disappeared and was replaced by a grey square of the same size. Participants were asked to maintain the outcome image in their minds as vividly as possible for another 3 s. Forty-eight pairs were presented randomly across the four encoding blocks. To ensure that participants fully encoded the pharmaceutical products-health outcomes associations before engaging in the counter-evaluative learning on the second day, only those who achieved an accuracy higher than the pre-defined criteria (80%) in the A-B cued recognition would proceed.

#### Counter-evaluative Learning Task (A-C association learning)

On the evening of Day 2, participants learned the counter-evaluative information about the pharmaceutical products. Specifically, they would learn the associations between the 32 previously studied pseudowords and other health outcome images that had the valence opposite to the Day 1 evening learning. The remaining 16 pseudowords served as filler pairs, such that these pseudowords were also presented but were not paired with new images. Similar to the evaluative learning task, the counter-evaluative learning task included four encoding blocks. To increase the authenticity, participants were given the following instructions at the beginning of the task:

*“Many pharmaceutical products have multifaceted effects: they may produce both positive and negative health outcomes simultaneously. In this task, you will learn about the additional health effects caused by the previously studied pharmaceutical products. Please note that, in some cases, no new information is obtained, and a mosaic will be displayed on the screen instead to indicate such a situation.”*

In each trial of the encoding blocks, a fixation was presented for 0.5 s, followed by a blank screen with a 0.8-1.2s jitter. Next, a previously studied pseudoword (i.e., hypothetical pharmaceutical product names) was aurally presented, together with a health outcome image with opposite valence or a mosaic image for the filler stimuli, being visually presented on the screen for 3 s. Participants were required to think and memorize the pseudoword-outcome associations during this encoding phase.

After that, the outcome image disappeared and was replaced by a grey square of the same size. Participants were asked to maintain the outcome image in their minds as vividly as possible for another 3 s. Forty-eight pairs were presented randomly across the four encoding blocks.

#### Speeded Choice Task

To quantify participants’ automatic and speeded evaluation of the hypothetical pharmaceutical products, participants were asked to decide whether they were willing to choose this item in a relatively short time (Xia, Yao, et al., 2023).

Participants completed the speeded choice task three times: after evaluative learning, after counter-evaluative learning, and after TMR. Each trial started with a 0.5 fixation, followed by a blank screen with a 0.8-1.2 s jitter. Next, a pseudoword was aurally presented, while participants were required to respond as soon as possible (“A” for yes, “L” for no) in 1.7 s. The speeded choice task contained three blocks, with 48 pseudowords randomly presented in each block. The percentage of choosing “Yes” was defined as “Choice Rate”.

#### Evaluative Classification with Mouse-tracking

Participants completed the evaluative classification task three times (**Figure 1B**): after Day 1 evening evaluative learning, after Day 2 evening counter-evaluative learning, and after the TMR on Day 3 morning. In the evaluative classification with the mouse-tracking task, each trial started with two response options (“Negative” or “Positive”) presented at each corner at the top of the screen for a jitter duration between 0.8-1.2 s. The position was randomized across all trials. Participants were asked to take a careful look at the positions of the two response options before the next step so that the interval between presenting the stimuli and moving the mouse could be shorter. Next, a start button was presented at the bottom of the screen so that participants would have to return to this common area before moving the mouse. After participants clicked on the start button, one of the 48 pseudowords was presented aurally, and in the meantime, participants were instructed to start moving the mouse toward the valence. This task set-up allows better tracking of participants’ mouse trajectories during decision making. Each trial ended when participants clicked on one of the two valence buttons. The streaming x- and y-coordinates were recorded. The evaluative classification task contained three blocks, with 48 pseudowords randomly presented in each block.

#### Explicit Healthiness Rating Task

To assess participants’ explicit evaluation of these hypothetical pharmaceutical products, we asked participants to evaluate the healthiness of all 48 pharmaceutical products two times: after the Day 1 evening evaluative learning and after the TMR on Day 3 morning. Note that participants did not complete the explicit rating task after the Day 2 evening counter-evaluative learning task to shorten the interval between the counter-evaluative learning and sleep/TMR. Each trial began with a 0.5 s fixation and a 0.8-1.2 s blank screen, followed by the aural presentation of the product names. Participants then evaluated the pharmaceutical product on the degree of health outcome resulting from it on a 1-9 scale (1 = Extremely bad, 11 = Extremely good), using a blue triangle presented on the screen.

#### Baseline A-B Mental Retrieval Task

To test whether there was a difference in memory between negative-to-positive and positive-to-negative valence reversals and between cued and uncued TMR conditions before the counter-evaluative learning task, participants completed the baseline mental retrieval task at the beginning of the Day 2 experiment. Each trial in the baseline mental retrieval task started with a 0.5 s fixation and was followed by a blank screen with a jitter duration between 0.8 to 1.2 s. Next, participants heard the pseudowords and were required to think about the associated images that they learned in the Day 1 evaluative learning task as vividly as possible. Afterward, participants reported whether they remembered the associated image (remembered or not) and judged the valence of the associated image (positive or negative) by pressing buttons on the keyboards for a maximum of 1.5 s. The baseline mental retrieval task contained three blocks, with 48 pseudowords randomly presented in each block.

#### A-B, A-C, and B-C Cued Recognition Task

We assessed participants’ recognition memory using the A-B, A-C, and B-C cued recognition tasks (**Figure 1C**). Each recognition task contained all 48 pairs, being presented randomly. Participants completed the A-B task after the evaluative learning task on the evening of Day 1, and the A-C task after the counter-evaluative learning task on the evening of Day 2. On the morning of Day 3 following the TMR, participants completed all three tasks (B-C, A-B, A-C), with the order of A-B and A-C being counter-balanced across participants. For a pseudoword, if participants correctly recognized the associated images in all three tasks (B-C/A-C/A-B), then this item would be coded as “integrated”. If participants made errors in either of the three tasks, the item would be coded as “non-integrated.”

In the A-B task, each trial began with a 0.5-s fixation followed by a blank screen with 0.8-1.2 s. Pseudowords were then auditorily presented, accompanied by a grey square at the screen’s center for 3 s. Participants were instructed to recall the associated health outcome images from the evaluative learning task as vividly as possible. They subsequently indicated within 1.5 s whether they remembered the associated images (i.e., subjective remembering) and then selected the correct image from four options within 3 s (i.e., cued recognition accuracy). To minimize familiarity effects, the three other images were chosen from the images that participants learned on the same day, with one sharing the same valence as the correct answer and the other two displaying opposite valence. Participants pressed the spacebar if they forgot the image.

Both the A-C and B-C tasks were similar to the A-B task procedure, with the following differences: In the A-C task, participants were asked to recall the associated health outcome images from the counter-evaluative learning task. In the B-C task, trials began with a 0.5-s fixation and a 0.8 to 1.2-s blank screen, followed by the visual presentation of the health outcome from the Day 1 evaluative learning task at the center of the screen for 3 ss. Participants were required to vividly recall the alternative health outcome images in the Day 2 evening counter-evaluative learning task, which were paired with the same pharmaceutical product (i.e., the pseudowords). In both A-C and B-C tasks, participants could press the spacebar to indicate if the pseudowords were paired with mosaic (i.e., no new information was provided in the counter-evaluative learning task).

### Day 2 NREM TMR

Half of the pseudowords from either the negative-to-positive or positive-to-negative conditions (16 out of 32, e.g., “Fajin”, memory cues) and 16 additional pseudowords (i.e., control cues) were played during the TMR (**Figure 1A**). These 16 pseudowords were presented during the pseudoword familiarization task, but were never paired with any images before sleep. Throughout the Day 2 night, pink noise was played as the background noise. Well-trained experimenters monitored the EEG brainwaves and identified the sleeping stages for TMR administration. For online sleep monitoring, F3/F4, C3/C4, P3/P4, O1/O2, electro-oculography (EOG), and electromyography (EMG), with online reference at CPz, were selected. Upon detection of stable slow-wave sleep for at least 5 minutes, the spoken pseudowords were played via a loudspeaker placed above the participant’s head. In each block of the TMR, all 32 cues were randomly played (∼1 s), followed by an inter-stimulus interval (ISI) of 6 ± 0.2 s. A 30-s interval separated each block of playing. The TMR was terminated when 20 cueing blocks were completed or when it reached 2 a.m., whichever came first. Cueing was paused immediately when participants showed signs of micro-arousal or awakening, or they entered N1 or REM sleep. Cueing would be resumed when participants returned to stable slow-wave sleep.

Participants were excluded if they received fewer than three TMR blocks (*n* = 5). We evaluated the accuracy of TMR cueing during NREM sleep, and found 97.51% (S.D. = 13.78%) of cues are played during NREM sleep.

### Mouse-tracking Quantification

The mouse-movement data was collected by *PsychoPy* (version 2020.1.3; Peirce et al., 2019), and were analyses with *MouseTrap* (version 3.2.1; Kieslich et al., 2018; Wulff et al., 2021) implemented in R (version 4.2.2). The mouse-tracking data were normalized temporally and spatially to facilitate comparison across trials and participants, considering the large differences in response times. Spatially, all trajectories were remapped to the left side of the coordinate system. Next, spatial normalization was employed by aligning the starting point across all trials and all subjects. Temporally, the trajectories were time-normalized by slicing the responses into 101 identical time bins using linear interpolation (Spivey & Dale, 2006).

Afterward, trials were excluded if the starting reaction times were longer than 0.625 s and the overall reaction times were longer than 5 s. Lastly, the mouse-tracking measures were calculated for each trajectory, such as different measures for curvature (**Figure 1B**), including maximum deviation (MAD), average deviation (AD), and the area under the curve (AUC), which would be an index of ambivalence level (Freeman & Ambady, 2010). We sum the MD, AD, and AUC together to have a comprehensive index of decision ambivalence (Xu et al., 2024).

### EEG Acquisition

Continuous EEGs were recorded with an eego amplifier and a 64-channel gel-based waveguard cap (10–20 layout, ANT Neuro, Enschede, and Netherlands). The online sampling rate was 500 Hz, with CPz as the online reference and AFz as the ground electrode. The horizontal electrooculogram (EOG) was recorded from an electrode placed 1.5 cm to the left external canthus. The vertical EOG was recorded from an electrode placed 1 cm below the left eye. Two additional electrodes were attached to both sides of the chins to measure electromyography (EMG). The impedance of all electrodes was maintained below 20 kΩ during the recording.

### EEG Preprocessing

Sleep EEG was processed offline using custom Python (3.8.8) scripts and MNE-Python (1.4.0; Gramfort et al., 2013). Unused channels (HEOG, VEOG, M1, and M2) were first removed from the EEG data. The raw EEG was then filtered with a bandpass filter of 0.5-40 Hz and was notch-filtered at 50 Hz. Afterward, the EEG was downsampled to 250 Hz to facilitate the following analyses. Bad channels were then visually detected, removed, and interpolated. The EEG data were next re-referenced to the whole-brain average, followed by segmentation into [-15 s to 15 s] epochs relative to the onset of the cue. Bad epochs were then visually detected and removed from further analyses. Artifact-free EEG data were further segmented into [-2 s to 7 s] epochs for time-frequency analysis. The number of remaining epochs for each condition is provided in Table S1. The overnight continuous EEG data were also retained for sleep staging and overnight spindle detection.

### Offline Automated Sleep Staging

The offline sleep staging was conducted with the YASA toolbox (0.6.3; Vallat & Walker, 2021) implemented in Python (3.8.8). Raw overnight continuous EEG data were re-referenced to M1 and M2 according to the YASA recommendation. Sleep staging was based on C4 (or C3 if C4 was marked as a bad channel), left horizontal EOG, and left EMG (see Table S2 for sleep stage information).

### Sleep Event Detection

The automated spindle detection was implemented in the YASA toolbox (0.6.3; Vallat & Walker, 2021). We applied two thresholds in identifying a spindle: correlation, the correlation between the sigma-filtered signal and broadband signal, and RMS, the moving root mean square (RMS) of the sigma-filtered signal. The spindle detection algorithm was applied to the artifact-free [-15 s to 15 s] epochs with a correlation of 0.50 and RMS of 1.5. Subsequently, the algorithm generated a series of binary values (spindle presence or absence) to indicate whether a spindle was detected at each timepoint (each timepoint represented 4 ms). The cue-elicited spindle probability was next determined by computing the proportion of detected spindles across trials at each timepoint (Schechtman et al., 2021; Xia, Yao, et al., 2023).

### Time-Frequency Analysis

For the time-frequency analysis, we focused on nine fronto-central channels (F1/2, Fz, FC1/2, FCz, C1/2, Cz) in accordance with recent studies examining auditory processing during sleep (Xia, Yao, et al., 2023; Züst et al., 2019). Morlet wavelets transformation with variance cycles (three cycles at 1 Hz in length, increasing linearly with frequency to 15 cycles at 30 Hz) was applied to the [-2 s to 7s] epochs to compute time-frequency representation (TFR) for the 1-30 Hz EEG. Next, epochs were further segmented into [-1s to 5s] epochs to eliminate edge artifacts. The trial-level spectral power was normalized (Z-scored) using [-1 s to -0.2 s] baseline of the averaged spectral power of all trials.

## Statistical Analysis

The impacts of counter-evaluative learning and TMR cueing on evaluation and the recognition memories were examined by conducting a series of 2 (valence reversals: negative-to-positive vs. positive-to-negative) by 2 (TMR conditions: cued vs. uncued) repeated measure ANOVAs.

We next assessed the impact of TMR on decision ambivalence. Because the number of valid items varied across participants and conditions, we used Bayesian item-level linear mixed-effects models (BLMMs), which can accommodate unbalanced data and explicitly model variability across both participants and items. Ambivalence derived from mouse-tracking was modeled as a function of TMR (cued vs. uncued), valence reversal (positive-to-negative vs. negative-to-positive), and Time (Day 2 pre-TMR vs. Day 3 post-TMR), including their interactions (Formula 1 in Table 1).

**Table 1.**
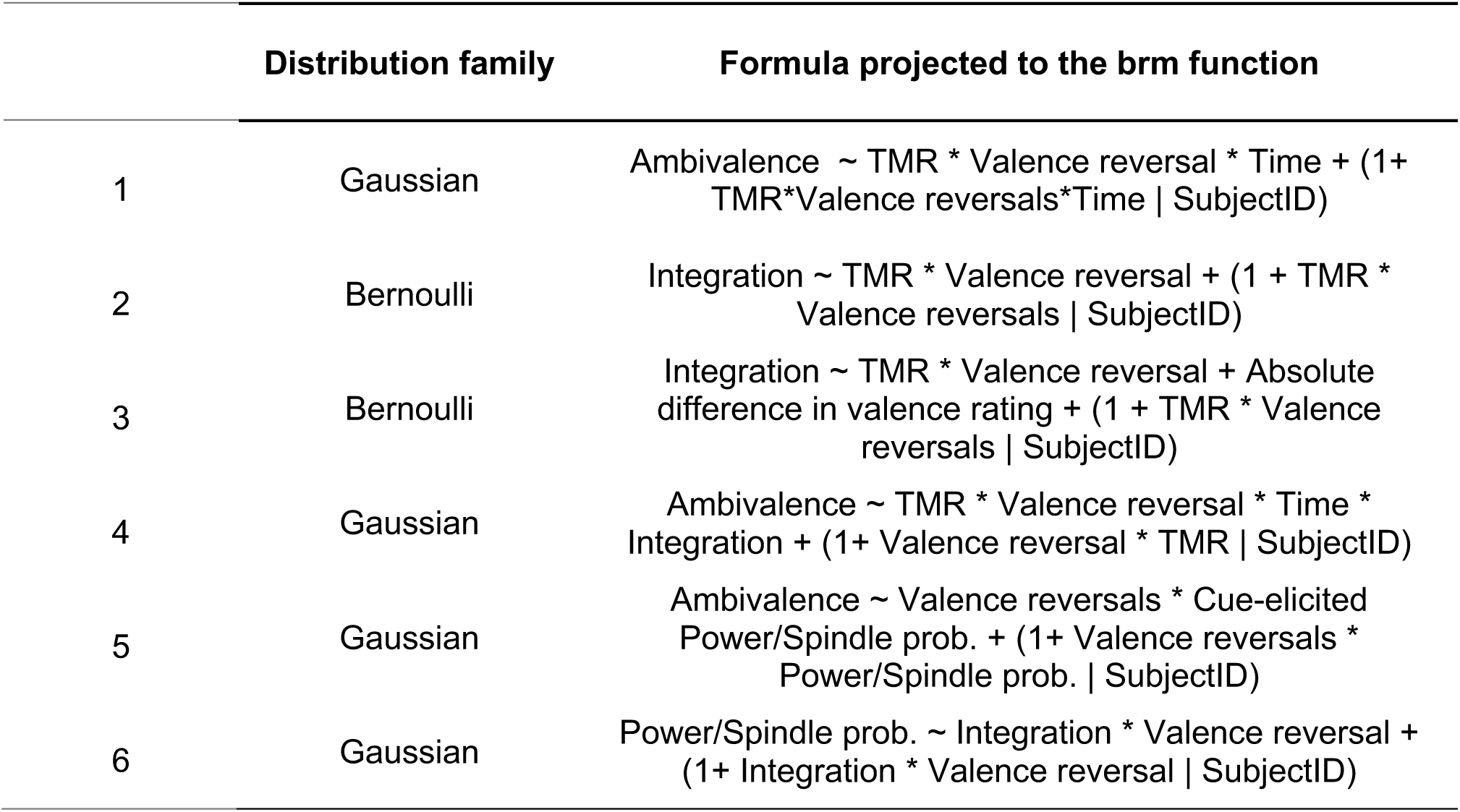
The Formula Applied in Bayesian Linear Mixed Model Analyses (BLMM)

To probe a memory mechanism, we tested whether TMR influenced evaluative memory integration (Day 2 pre-TMR and Day 3 post-TMR). Because the integration outcome was binary (integrated vs. non-integrated), we fit a Bayesian mixed-effects logistic model with a Bernoulli distribution (Formula 2 in Table 1). In addition, we included the absolute difference in healthiness ratings between the positive and negative images paired with the same pseudoword as a covariate to control for baseline evaluative disparity within each pair (Formula 3 in Table 1).

Taken together, to test whether memory integration was related to ambivalence reduction, we fit a BLMM including TMR, valence reversal, Time, and integration status (and their interactions) as fixed effects (Formula 4 in Table 1).

To characterize brain–behavior relationships, we first identified cue-elicited EEG responses using cluster-based permutation tests on (i) time-resolved spindle probability and (ii) time–frequency–resolved EEG power. We used two-tailed one-sample cluster permutation tests (MNE-Python), with 1,000 permutations and a cluster-forming threshold of 𝛼 = .05. We then extracted, for each participant, the mean cue-elicited delta power and spindle probability within the significant above-baseline time (and time–frequency) windows for subsequent brain–behavior analyses. Next, we tested brain–behavior associations using BLMMs. To relate cue-elicited delta power and spindle probability to ambivalence, we fit models including the cue-elicited activities (delta power or spindle probability), valence reversal, and their interaction as fixed effects (Formula 5 in Table 1). To relate the same EEG measures to memory integration, we fit BLMMs predicting integration status from the EEG metric, valence reversal, and their interaction (Formula 6 in Table 1).

All the repeated-measure ANOVAs were conducted with the *afex* package (1.2.1) implemented in R. All the BLMMs were conducted with the brms package (2.20.4; Bürkner, 2021) implemented in R. All post-hoc analyses were conducted with the emmeans package (1.8.7). All the formulas projected to the brm function was provided in **Table 1**. Statistical inferences for the BLMM were based on the 95% highest density interval (HDI) of the posterior distribution. Effects were considered credible if the 95% HDI did not encompass 0.

Finally, we investigated whether memory vs. control cues would elicit significantly different EEG power changes and spindle probability. We employed a cluster-based two-tailed one-sample permutation test, implemented in the *MNE-Python* toolbox with 1000 randomizations and a statistical threshold of 0.05.

## Acknowledgment

We thank Ruoying Zheng for her assistance in data collection. The research was supported by the Ministry of Science and Technology of China, STI2030-Major Projects (No. 2022ZD0214100), the National Natural Science Foundation of China (No. 32171056) to X.H.

## Author contribution

D.C.: conceptualization, investigation, formal analysis, data curation, software, methodology, writing – original draft, writing – review & editing, and visualization; T.X.: conceptualization, methodology, and writing –review & editing; J.L.: conceptualization and writing –review & editing; Y.Z. and X.Z.: investigation and writing – review & editing; H.W.: methodology, and writing-review & editing; C.S.W.L.: writing – review & editing, and funding acquisition; X.H.: conceptualization, writing - original draft, writing - review & editing, supervision, project administration, and funding acquisition.

## Data and code availability

Pre-processed data and statistical analysis scripts will be made available on the Open Science Framework (OSF) upon publication.

## Reference

1. Abdellahi, M. E. A., Rakowska, M., Treder, M. S., & Lewis, P. A. (2026). Targeted memory reactivation elicits temporally compressed reactivation linked to spindles. Imaging Neuroscience, 4, IMAG.a.1123. 10.1162/IMAG.a.1123

2. Abdellahi, M. E., Koopman, A. C., Treder, M. S., & Lewis, P. A. (2023). Targeted memory reactivation in human REM sleep elicits detectable reactivation. eLife, 12, e84324. 10.7554/eLife.84324

3. Ai, S., Yin, Y., Chen, Y., Wang, C., Sun, Y., Tang, X., Lu, L., Zhu, L., & Shi, J. (2018). Promoting subjective preferences in simple economic choices during nap. eLife, 7, e40583. 10.7554/eLife.40583

4. Antony, J. W., & Schechtman, E. (2023). Reap while you sleep: Consolidation of memories differs by how they were sown. Hippocampus, hipo.23526. 10.1002/hipo.23526

5. Antony, J. W., Schönauer, M., Staresina, B. P., & Cairney, S. A. (2019). Sleep Spindles and Memory Reprocessing. Trends in Neurosciences, 42(1), 1–3. 10.1016/j.tins.2018.09.012

6. Ashton, J. E., Cairney, S. A., & Gaskell, M. G. (2018). No effect of targeted memory reactivation during slow-wave sleep on emotional recognition memory. Journal of Sleep Research, 27(1), 129–137. 10.1111/jsr.12542

7. Béna, J., Mauclet, A., & Corneille, O. (2022). Does co-occurrence information influence evaluations beyond relational meaning? An investigation using self-reported and mouse-tracking measures of attitudinal ambivalence. Journal of Experimental Psychology: General. 10.1037/xge0001308

8. Biderman, N., Bakkour, A., & Shohamy, D. (2020). What Are Memories For? The Hippocampus Bridges Past Experience with Future Decisions. Trends in Cognitive Sciences, 24(7), 542–556. 10.1016/j.tics.2020.04.004

9. Biderman, N., Gershman, S. J., & Shohamy, D. (2023). The role of memory in counterfactual valuation. Journal of Experimental Psychology: General, 152(6), 1754–1767. 10.1037/xge0001364

10. Biderman, N., & Shohamy, D. (2021). Memory and decision making interact to shape the value of unchosen options. Nature Communications, 12(1), 4648. 10.1038/s41467-021-24907-x

11. Brodt, S., Inostroza, M., Niethard, N., & Born, J. (2023). Sleep—A brain-state serving systems memory consolidation. Neuron, 111(7), 1050–1075. 10.1016/j.neuron.2023.03.005

12. Bürkner, P.-C. (2021). Bayesian Item Response Modeling in R with brms and Stan. Journal of Statistical Software, 100, 1–54. 10.18637/jss.v100.i05

13. Cai, D. J., Mednick, S. A., Harrison, E. M., Kanady, J. C., & Mednick, S. C. (2009). REM, not incubation, improves creativity by priming associative networks. Proceedings of the National Academy of Sciences, 106(25), 10130–10134. 10.1073/pnas.0900271106

14. Cairney, S. A., Guttesen, A. á V., El Marj, N., & Staresina, B. P. (2018). Memory Consolidation Is Linked to Spindle-Mediated Information Processing during Sleep. Current Biology, 28(6), 948–954.e4. 10.1016/j.cub.2018.01.087

15. Cairney, S. A., Lindsay, S., Sobczak, J. M., Paller, K. A., & Gaskell, M. G. (2016). The Benefits of Targeted Memory Reactivation for Consolidation in Sleep are Contingent on Memory Accuracy and Direct Cue-Memory Associations. Sleep, 39(5), 1139–1150. 10.5665/sleep.5772

16. Chen, D., Xia, T., Yao, Z., Zhang, L., & Hu, X. (2024). Modulating social learning-induced evaluation updating during human sleep. Npj Science of Learning, 9(1), 1–11. 10.1038/s41539-024-00255-5

17. Cho, M., Murugavel, S., Thiha, A. S., & Schechtman, E. (2025). The effects of targeted reactivation on memories cued once or multiple times during a nap. Neuropsychologia, 219, 109275. 10.1016/j.neuropsychologia.2025.109275

18. Cowan, E., Liu, A., Henin, S., Kothare, S., Devinsky, O., & Davachi, L. (2020). Sleep Spindles Promote the Restructuring of Memory Representations in Ventromedial Prefrontal Cortex through Enhanced Hippocampal–Cortical Functional Connectivity. The Journal of Neuroscience, 40(9), 1909–1919. 10.1523/JNEUROSCI.1946-19.2020

19. Cox, W. R., Dobbelaar, S., Meeter, M., Kindt, M., & van Ast, V. A. (2021). Episodic memory enhancement versus impairment is determined by contextual similarity across events. Proceedings of the National Academy of Sciences, 118(48), e2101509118. 10.1073/pnas.2101509118

20. Creery, J. D., Oudiette, D., Antony, J. W., & Paller, K. A. (2015). Targeted Memory Reactivation during Sleep Depends on Prior Learning. Sleep, 38(5), 755–763. 10.5665/sleep.4670

21. del Río, M., Trudel, N., Prabhu, G., Hunt, L. T., Moutoussis, M., Dolan, R. J., & Hauser, T. U. (2026). Indecision and recency-weighted evidence integration in non-clinical and clinical settings. Nature Human Behaviour, 1–14. 10.1038/s41562-025-02385-1

22. Dudai, Y., Karni, A., & Born, J. (2015). The Consolidation and Transformation of Memory. Neuron, 88(1), 20–32. 10.1016/j.neuron.2015.09.004

23. Durrant, S. J., Taylor, C., Cairney, S., & Lewis, P. A. (2011). Sleep-dependent consolidation of statistical learning. Neuropsychologia, 49(5), 1322–1331. 10.1016/j.neuropsychologia.2011.02.015

24. Eil, D., & Rao, J. M. (2011). The Good News-Bad News Effect: Asymmetric Processing of Objective Information about Yourself. American Economic Journal. Microeconomics, 3(2), 114–138. 10.1257/mic.3.2.114

25. Ellenbogen, J. M., Hu, P. T., Payne, J. D., Titone, D., & Walker, M. P. (2007). Human relational memory requires time and sleep. Proceedings of the National Academy of Sciences, 104(18), 7723–7728. 10.1073/pnas.0700094104

26. Fernandez, L. M. J., & Lüthi, A. (2020). Sleep Spindles: Mechanisms and Functions. Physiological Reviews, 100(2), 805–868. 10.1152/physrev.00042.2018

27. Foster, D. W., Neighbors, C., & Prokhorov, A. (2014). Drinking motives as moderators of the effect of ambivalence on drinking and alcohol-related problems. Addictive Behaviors, 39(1), 133–139. 10.1016/j.addbeh.2013.09.016

28. Freeman, J. B., & Ambady, N. (2010). MouseTracker: Software for studying real-time mental processing using a computer mouse-tracking method. Behavior Research Methods, 42(1), 226–241. 10.3758/BRM.42.1.226

29. Gramfort, A., Luessi, M., Larson, E., Engemann, D., Strohmeier, D., Brodbeck, C., Goj, R., Jas, M., Brooks, T., Parkkonen, L., & Hämäläinen, M. (2013). MEG and EEG data analysis with MNE-Python. Frontiers in Neuroscience, 7. https://www.frontiersin.org/articles/10.3389/fnins.2013.00267

30. Hennies, N., Lambon Ralph, M. A., Kempkes, M., Cousins, J. N., & Lewis, P. A. (2016). Sleep Spindle Density Predicts the Effect of Prior Knowledge on Memory Consolidation. The Journal of Neuroscience, 36(13), 3799–3810. 10.1523/JNEUROSCI.3162-15.2016

31. Heycke, T., & Gawronski, B. (2020). Co-occurrence and relational information in evaluative learning: A multinomial modeling approach. Journal of Experimental Psychology: General, 149(1), 104–124. 10.1037/xge0000620

32. Hu, X., Cheng, L. Y., Chiu, M. H., & Paller, K. A. (2020). Promoting memory consolidation during sleep: A meta-analysis of targeted memory reactivation. Psychological Bulletin, 146(3), 218–244. 10.1037/bul0000223

33. Jin, R., Xia, T., Gawronski, B., & Hu, X. (2023). Attitudinal Effects of Stimulus Co-occurrence and Stimulus Relations: Sleep Supports Propositional Learning Via Memory Consolidation. Social Psychological and Personality Science, 14(1), 51–59. 10.1177/19485506211067673

34. Kieslich, P. J., Henninger, F., Wulff, D. U., Haslbeck, J. M. B., & Schulte-Mecklenbeck, M. (2018). Mouse-tracking: A practical guide to implementation and analysis [Preprint]. PsyArXiv. https://osf.io/zuvqa

35. Kim, S., Pjesivac, I., & Jin, Y. (2019). Effects of Message Framing on Influenza Vaccination: Understanding the Role of Risk Disclosure, Perceived Vaccine Efficacy, and Felt Ambivalence. Health Communication, 34(1), 21–30. 10.1080/10410236.2017.1384353

36. Küpper, C. S., Benoit, R. G., Dalgleish, T., & Anderson, M. C. (2014). Direct suppression as a mechanism for controlling unpleasant memories in daily life. Journal of Experimental Psychology: General, 143(4), 1443. 10.1037/a0036518

37. Lang, P. J., Bradley, M. M., & Cuthbert, B. N. (2005). International affective picture system (IAPS): Affective ratings of pictures and instruction manual. NIMH, Center for the Study of Emotion & Attention Gainesville, FL.

38. Lewis, P. A., & Durrant, S. J. (2011). Overlapping memory replay during sleep builds cognitive schemata. Trends in Cognitive Sciences, 15(8), 343–351. 10.1016/j.tics.2011.06.004

39. Lewis, P. A., Knoblich, G., & Poe, G. (2018). How Memory Replay in Sleep Boosts Creative Problem-Solving. Trends in Cognitive Sciences, 22(6), 491–503. 10.1016/j.tics.2018.03.009

40. Lim, S.-L., Penrod, M. T., Ha, O.-R., Bruce, J. M., & Bruce, A. S. (2018). Calorie Labeling Promotes Dietary Self-Control by Shifting the Temporal Dynamics of Health- and Taste-Attribute Integration in Overweight Individuals. Psychological Science, 29(3), 447–462. 10.1177/0956797617737871

41. Liu, J., Chen, D., Xia, T., Zeng, S., Xue, G., & Hu, X. (2025). Slow-wave sleep and REM sleep differentially contribute to memory representational transformation. Communications Biology, 8, 1407. 10.1101/2024.08.05.606592

42. Liu, J., Xia, T., Chen, D., Yao, Z., Zhu, M., Antony, J. W., Lee, T. M. C., & Hu, X. (2023). Item-specific neural representations during human sleep support long-term memory. PLOS Biology, 21(11), e3002399. 10.1371/journal.pbio.3002399

43. Luo, Y. J., & Wang, Y. N. (2004). Chinese affective words system (CAWS). Beijing: Institute of Psychology, CAS.

44. Melnikoff, D. E., Mann, T. C., Stillman, P. E., Shen, X., & Ferguson, M. J. (2021). Tracking Prejudice: A Mouse-Tracking Measure of Evaluative Conflict Predicts Discriminatory Behavior. Social Psychological and Personality Science, 12(2), 266–272. 10.1177/1948550619900574

45. Menninga, K. M., Dijkstra, A., & Gebhardt, W. A. (2011). Mixed feelings: Ambivalence as a predictor of relapse in ex-smokers. British Journal of Health Psychology, 16(3), 580– 591. 10.1348/135910710X533219

46. Nicholas, J., Daw, N. D., & Shohamy, D. (2025). Proactive and reactive construction of memory-based preferences. Nature Communications, 16(1), 1618. 10.1038/s41467-025-56183-4

47. Oser, M. L., McKellar, J., Moos, B. S., & Moos, R. H. (2010). Changes in ambivalence mediate the relation between entering treatment and change in alcohol use and problems. Addictive Behaviors, 35(4), 367–369. 10.1016/j.addbeh.2009.10.024

48. Ou, J., Qu, Y., Xu, Y., Xiao, Z., Behrens, T., & Liu, Y. (2025). Replay builds an efficient cognitive map offline to avoid computation online. Neuroscience. 10.1101/2025.01.08.632067

49. Oudiette, D., & Paller, K. A. (2013). Upgrading the sleeping brain with targeted memory reactivation. Trends in Cognitive Sciences, 17(3), 142–149. 10.1016/j.tics.2013.01.006

50. Paller, K. A., Creery, J. D., & Schechtman, E. (2021). Memory and Sleep: How Sleep Cognition Can Change the Waking Mind for the Better. Annual Review of Psychology, 72(1), 123–150. 10.1146/annurev-psych-010419-050815

51. Peirce, J., Gray, J. R., Simpson, S., MacAskill, M., Höchenberger, R., Sogo, H., Kastman, E., & Lindeløv, J. K. (2019). PsychoPy2: Experiments in behavior made easy. Behavior Research Methods, 51(1), 195–203.

52. Pereira, S. I. R., Santamaria, L., Andrews, R., Schmidt, E., Van Rossum, M., & Lewis, P. A. (2023). Rule abstraction is facilitated by auditory cueing in REM sleep. *The Journal of Neuroscience*, JN-RM-1966–21. 10.1523/JNEUROSCI.1966-21.2022

53. Rakowska, M., Abdellahi, M. E. A., Bagrowska, P., Navarrete, M., & Lewis, P. A. (2021). Long term effects of cueing procedural memory reactivation during NREM sleep. NeuroImage, 244, 118573. 10.1016/j.neuroimage.2021.118573

54. Rasch, B., Büchel, C., Gais, S., & Born, J. (2007). Odor Cues During Slow-Wave Sleep Prompt Declarative Memory Consolidation. Science, 315(5817), 1426–1429. 10.1126/science.1138581

55. Rihm, J. S., Diekelmann, S., Born, J., & Rasch, B. (2014). Reactivating Memories during Sleep by Odors: Odor Specificity and Associated Changes in Sleep Oscillations. Journal of Cognitive Neuroscience, 26(8), 1806–1818. 10.1162/jocn_a_00579

56. Rudoy, J. D., Voss, J. L., Westerberg, C. E., & Paller, K. A. (2009). Strengthening Individual Memories by Reactivating Them During Sleep. Science, 326(5956), 1079–1079. 10.1126/science.1179013

57. Schechtman, E., Antony, J. W., Lampe, A., Wilson, B. J., Norman, K. A., & Paller, K. A. (2021). Multiple memories can be simultaneously reactivated during sleep as effectively as a single memory. Communications Biology, 4(1), 25. 10.1038/s42003-020-01512-0

58. Schechtman, E., Heilberg, J., & Paller, K. A. (2023). Memory consolidation during sleep involves context reinstatement in humans. Cell Reports, 42(4), 112331. 10.1016/j.celrep.2023.112331

59. Schneider, I. K., & Schwarz, N. (2017). Mixed feelings: The case of ambivalence. *Current Opinion in Behavioral Sciences*, Mixed Emotions, 15, 39–45. 10.1016/j.cobeha.2017.05.012

60. Sharot, T., & Garrett, N. (2016). Forming Beliefs: Why Valence Matters. Trends in Cognitive Sciences, 20(1), 25–33. 10.1016/j.tics.2015.11.002

61. Shohamy, D., & Daw, N. D. (2015). Integrating memories to guide decisions. *Current Opinion in Behavioral Sciences*, Neuroeconomics, 5, 85–90. 10.1016/j.cobeha.2015.08.010

62. Siefert, E. M., Uppuluri, S., Mu, J., Tandoc, M. C., Antony, J. W., & Schapiro, A. C. (2024). Memory Reactivation during Sleep Does Not Act Holistically on Object Memory. Journal of Neuroscience, 44(24). 10.1523/JNEUROSCI.0022-24.2024

63. Son, J.-Y., Vives, M.-L., Bhandari, A., & FeldmanHall, O. (2024). Replay shapes abstract cognitive maps for efficient social navigation. Nature Human Behaviour. 10.1038/s41562-024-01990-w

64. Spivey, M. J., & Dale, R. (2006). Continuous Dynamics in Real-Time Cognition. Current Directions in Psychological Science, 15(5), 207–211. 10.1111/j.1467-8721.2006.00437.x

65. Tamminen, J., Lambon Ralph, M. A., & Lewis, P. A. (2017). Targeted memory reactivation of newly learned words during sleep triggers REM-mediated integration of new memories and existing knowledge. Neurobiology of Learning and Memory, 137, 77– 82. 10.1016/j.nlm.2016.11.012

66. Tamminen, J., Payne, J. D., Stickgold, R., Wamsley, E. J., & Gaskell, M. G. (2010). Sleep Spindle Activity is Associated with the Integration of New Memories and Existing Knowledge. Journal of Neuroscience, 30(43), 14356–14360. 10.1523/JNEUROSCI.3028-10.2010

67. Temudo, A., & Albouy, G. (2024). Using targeted memory reactivation as a tool to provide mechanistic insights into memory consolidation during sleep. *Sleep*, zsae163. 10.1093/sleep/zsae163

68. Vallat, R., & Walker, M. P. (2021). An open-source, high-performance tool for automated sleep staging. eLife, 10, e70092. 10.7554/eLife.70092

69. van Harreveld, F., van der Pligt, J., & de Liver, Y. N. (2009). The Agony of Ambivalence and Ways to Resolve It: Introducing the MAID Model. Personality and Social Psychology Review, 13(1), 45–61. 10.1177/1088868308324518

70. van Kesteren, M. T. R., Krabbendam, L., & Meeter, M. (2018). Integrating educational knowledge: Reactivation of prior knowledge during educational learning enhances memory integration. Npj Science of Learning, 3(1), 1–8. 10.1038/s41539-018-0027-8

71. Wang, J., Otgaar, H., Smeets, T., Howe, M. L., & Zhou, C. (2019). Manipulating memory associations changes decision-making preferences in a preconditioning task. Consciousness and Cognition, 69, 103–112. 10.1016/j.concog.2019.01.016

72. Wimmer, G. E., & Shohamy, D. (2012). Preference by Association: How Memory Mechanisms in the Hippocampus Bias Decisions. Science, 338(6104), 270–273. 10.1126/science.1223252

73. Wulff, D. U., Kieslich, P. J., Henninger, F., Haslbeck, J., & Schulte-Mecklenbeck, M. (2021). Movement tracking of cognitive processes: A tutorial using mousetrap. PsyArXiv. 10.31234/osf.io/v685r

74. Xia, T., Antony, J. W., Paller, K. A., & Hu, X. (2023). Targeted memory reactivation during sleep influences social bias as a function of slow-oscillation phase and delta power. Psychophysiology, 60(5), e14224. 10.1111/psyp.14224

75. Xia, T., Chen, D., Zeng, S., Yao, Z., Liu, J., Qin, S., Paller, K. A., Torres Platas, S. G., Antony, J. W., & Hu, X. (2024). Aversive memories can be weakened during human sleep via the reactivation of positive interfering memories. Proceedings of the National Academy of Sciences, 121(31), e2400678121. 10.1073/pnas.2400678121

76. Xia, T., & Hu, X. (2025). Memory editing during sleep: Mechanisms, clinical applications, and technological innovations. Trends in Cognitive Sciences, 0(0). 10.1016/j.tics.2025.07.010

77. Xia, T., Yao, Z., Guo, X., Liu, J., Chen, D., Liu, Q., Paller, K. A., & Hu, X. (2023). Updating memories of unwanted emotions during human sleep. Current Biology, 33(2), 309–320.e5. 10.1016/j.cub.2022.12.004

78. Xu, X. J., Liu, X., Hu, X., & Wu, H. (2025). The trajectory of crime: Integrating mouse-tracking into concealed memory detection. Behavior Research Methods, 57(2), 78. 10.3758/s13428-024-02594-y

79. Xu, X. J., Mobbs, D., & Wu, H. (2024). Unethical amnesia brain: Memory and metacognitive distortion induced by dishonesty. Neuroscience. 10.1101/2024.03.03.583239

80. Yao, Z., Lin, X., & Hu, X. (2021). Optimistic amnesia: How online and offline processing shape belief updating and memory biases in immediate and long-term optimism biases. Social Cognitive and Affective Neuroscience, 16(5), 453–462. 10.1093/scan/nsab011

81. Yuksel, C., Denis, D., Coleman, J., Ren, B., Oh, A., Cox, R., Morgan, A., Sato, E., & Stickgold, R. (2025). Both slow wave and rapid eye movement sleep contribute to emotional memory consolidation. Communications Biology, 8(1), 485. 10.1038/s42003-025-07868-5

82. Zarocostas, J. (2020). How to fight an infodemic. The Lancet, 395(10225), 676. 10.1016/S0140-6736(20)30461-X

83. Züst, M. A., Ruch, S., Wiest, R., & Henke, K. (2019). Implicit Vocabulary Learning during Sleep Is Bound to Slow-Wave Peaks. Current Biology, 29(4), 541–553.e7. 10.1016/j.cub.2018.12.038

