## Supplementary material for "Reactivating conflicting evaluative memories during sleep reduces decision ambivalence": Data S1, Data S2, Data S3, Data S4, Data S5, Data S6, Table S1, Table S2

### Data S1. Effective Evaluative Learning on Memory and Evaluation

We first examined how evaluative learning influenced evaluative memories (A-B cued recognition) using the 2 (valence reversals: negative-to-positive vs. positive-to-negative) by 2 (TMR: cued vs. uncued) repeated measure ANOVAs. Results showed that participants fully encoded the evaluative learning: The A-B recognition accuracy was high (Mean ± S.D., 0.94 ± 0.09) and showed no difference related to TMR and valence reversals (*p*s > .169).

The successful evaluative learning further shaped participants' evaluations effectively: Positive evaluative learning led to higher choice rate in the speeded choice task ($F$ (1, 35) = 167.11, $p$ < .001, $\eta_G^2$ = 0.650), higher positivity classification in the evaluative classification task ($F$ (1, 35) = 173.92, $p$ < .001, $\eta_G^2$ = 0.728), and higher healthiness ratings than negative learning ($F$ (1, 35) = 284.96, $p$ < .001, $\eta_G^2$ = 0.814). In addition, neither the main effect of TMR nor their interaction with valence reversals was significant (*p*s > .154).

These results justified that the evaluative learning (pairing hypothetical pharmaceutical products and the valanced health outcomes) successfully formed initial evaluations, as evidenced by high memory accuracy and significant changes in evaluations.

### Data S2. Effective Counter-evaluative Learning on Memory and Evaluation

We next examined how counter-evaluative learning influenced counter-evaluative memories (A-C cued recognition) and evaluations, using the 2 (valence reversals: negative-to-positive vs. positive-to-negative) by 2 (TMR: cued vs. uncued) repeated measure ANOVA. Results showed that participants fully successfully encoded the counter-evaluative learning: The A-C recognition accuracy was high (Mean ± S.D., 0.84 ± 0.18). However, we unexpectedly found a significant main effect of valence reversals on the A-C cued recognition accuracy (negative-to-positive > positive-to-negative, $F$ (1, 35) = 6.44, $p$ = .016, $\eta_G^2$ = .029). Neither TMR nor their interaction effect was found (*p*s > .547).

The same valence reversals by TMR repeated measure ANOVA showed that the successful counter-evaluative learning further updated participants' evaluation: There were significantly higher changes of the negative-to-positive than the positive-to-negative conditions in choice rate ($F$ (1, 34) = 57.86, $p$ < .001, $\eta_G^2$ = .472), and positivity rate ($F$ (1, 35) = 64.08, $p$ < .001, $\eta_G^2$ = .554), from Day 1 post-evaluative learning to

Day 2 post-counter-evaluative learning sessions. Neither the main effects of TMR nor their interaction effects were found ($p$s > .176).

These results revealed that learning counter-evaluative information updated the evaluation and confirmed that the evaluation of cued and uncued items was comparable before TMR cueing.

**Data S3. TMR effects on Mouse-tracking Trajectories**

We fit item-level Bayesian linear mixed-effects models (BLMMs) predicting mouse-tracking ambivalence indices from TMR (cued vs. uncued), valence reversal (negative-to-positive vs. positive-to-negative), and Time (Day 2 post-counter-evaluative learning vs. Day 3 post-TMR), including their interactions.

For Area Under Curve (AUC), the model showed a significant TMR by session interaction ($Median_{diff}$ = 0.08, 95% HDI [0.03, 0.14]). Post-hoc analyses indicated that cueing reduced ambivalence from Day 2 post-counter-evaluative learning to post-TMR (post-counter-evaluative learning vs. post-TMR, $Median_{diff}$ = 0.04, 95% HDI [0.01, 0.07]), while uncued items showed no significant changes ($Median_{diff}$ = -0.01, 95% HDI [-0.04, 0.02]).

For Area Under Curve (AUC), the model showed a significant TMR by session interaction ($Median_{diff}$ = 0.08, 95% HDI [0.03, 0.14]). Post-hoc analyses indicated that cueing reduced ambivalence from Day 2 post-counter-evaluative learning to post-TMR (post-counter-evaluative learning vs. post-TMR, $Median_{diff}$ = 0.04, 95% HDI [0.01, 0.07]), while uncued items showed no significant changes ($Median_{diff}$ = -0.01, 95% HDI [-0.04, 0.02]).

For Maximum Deviation (MD), the model again showed a significant TMR by session interaction ($Median_{diff}$ = 0.08, 95% HDI [0.01, 0.15]). Post-hoc analyses revealed significant cueing numerically reduced ambivalence from Day 2 post-counter-evaluative learning and post-TMR (post-counter-evaluative learning vs. post-TMR, $Median_{diff}$ = 0.04, 95% HDI [-0.01, 0.08]), while uncued items showed no significant changes ($Median_{diff}$ = -0.02, 95% HDI [-0.06, 0.03]).

For Average Deviation (AD), the model again did not show a significant TMR by session interaction ($Median_{diff}$ = 0.02, 95% HDI [-0.00, 0.05]).

**Data S4. TMR effects on Evaluations**

We next tested whether TMR cueing further changed participants' evaluations after sleep. We examined the choice rate in the speeded choice task, and the positivity rate (i.e., choosing "positive") in the evaluative classification task. We treated performance on the Day 2 post-counter-evaluative learning session as the baseline and calculated change scores from Day 2 post-counter-evaluative learning to Day 3 post-TMR. A repeated-measures ANOVA with valence reversal (negative-to-positive vs. positive-to-negative) and TMR (cued vs. uncued) showed no reliable effects on changes in either choice rate or positivity rate ($p$s > .270).

For explicit healthiness ratings, because ratings were not collected on the Day 2 night, we used the Day 1 post-evaluative learning ratings as baseline and computed change scores from Day 1 post-evaluative learning to Day 3 post-TMR. The same valence reversal by TMR ANOVA on rating changes showed a strong main effect of valence reversal: ratings shifted more positively in the negative-to-positive condition than in the positive-to-negative condition ($F(1, 35) = 95.87$, $p < .001$, $\eta_p^2 = .653$). However, there was no evidence that TMR further changed ratings, as neither the main effect of TMR nor the interaction was significant (*p*s > .336).

Overall, though counter-evaluative learning updated evaluations and this shift was still present after one night of sleep, TMR did not further augment evaluation updating.

**Data S5. TMR Effects on AB/AC/BC Cued Recognition Tests**

We next examined whether TMR cueing impacted the performance in the memory tests respectively. To this end, we fitted item-level Bayesian linear mixed-effects models (BLMMs) predicting A-B, A-C, B-C cued recognition tests performance from TMR (cued vs. uncued), and valence reversal (negative-to-positive vs. positive-to-negative), including their interactions as fix factors.

For A-B cued recognition tests, no significant main effect of valence reversal ($Median_{diff}$ = -0.48, 95% HDI [-1.15, 1.13]), TMR ($Median_{diff}$ = -0.15, 95% HDI [-0.89, 0.58]) and their interaction ($Median_{diff}$ = 0.85, 95% HDI [-0.13, 1.84]) were found.

For A-C cued recognition tests, no significant main effect of valence reversal ($Median_{diff}$ = -0.40, 95% HDI [-0.90, 0.08]), TMR ($Median_{diff}$ = -0.10, 95% HDI [-0.64, 0.44]) and their interaction ($Median_{diff}$ = 0.21, 95% HDI [-0.51, 0.85]) were found.

For B-C cued recognition tests, no significant main effect of valence reversal ($Median_{diff}$ = -0.27, 95% HDI [-0.68, 0.13]), TMR ($Median_{diff}$ = -0.05, 95% HDI [-0.44, 0.35]) and their interaction ($Median_{diff}$ = 0.34, 95% HDI [-0.22, 0.90]) were found.

Together, these results showed no significant effects of TMR on single memory tests.

**Data S6. TMR effects on A-B-C Integration Controlling valence difference**

To control the valence difference at the item level, we fitted a BLMM with TMR and valence reversals as fix factors, and absolute difference in healthiness ratings at item levels as a covariate. We again found a significant main effect of valence reversal (positive-to-negative vs. negative-to-positive; $Median_{diff}$ = -0.47, 95% HDI [-0.89, -0.10]), and a significant TMR by valence reversal interaction effect ($Median_{diff}$ = 0.61, 95% HDI [0.07, 1.15]). Consistent with the models without covariates, post-hoc analyses revealed that, among cued items, integration was higher in the negative-to-positive than the positive-to-negative condition (cued negative-to-positive vs. cued positive-to-negative, $Median_{diff}$ = 0.47, 95% HDI [0.10, 0.89]), whereas this valence reversal difference was not significant among uncued items (uncued negative-to-positive vs. uncued positive-to-negative, $Median_{diff}$ = -0.14, 95% HDI [-0.54, 0.27]).

Further, within the positive-to-negative condition, TMR cueing was associated with reduced integration (cued vs. uncued, $Median_{diff}$ = -0.42, 95% HDI [-0.80, -0.01]).

**Table S1**

*Number of Artifact-Free Epochs in the TMR*

| Conditions | Mean | S.D. |
|---|---|---|
| **Negative-to-Positive** | 89.24 | 31.81 |
| **Positive-to-Negative** | 89.17 | 31.97 |
| **Control** | 178.57 | 64.05 |
| **In total** | 356.98 | 127.79 |

**Table S2**

*Sleep Staging Statistics*

| Item | Mean | S.D. |
|---|---|---|
| **TIB** | 481.07 | 7.06 |
| **SPT** | 454.95 | 31.79 |
| **WASO** | 44.68 | 59.60 |
| **TST** | 410.27 | 73.09 |
| **N1** | 30.94 | 14.08 |
| **N2** | 215.96 | 37.08 |
| **N3** | 76.67 | 23.43 |
| **REM** | 86.70 | 28.42 |
| **NREM** | 323.57 | 53.67 |
| **SOL** | 20.94 | 23.34 |
| **Lat_N1** | 24.27 | 31.06 |
| **Lat_N2** | 24.43 | 23.27 |
| **Lat_N3** | 34.26 | 22.37 |
| **Lat_REM** | 101.04 | 48.67 |
| **%N1** | 7.38 | 2.91 |
| **%N2** | 53.57 | 8.98 |
| **%N3** | 18.60 | 5.71 |
| **%REM** | 20.45 | 5.84 |
| **%NREM** | 79.55 | 5.84 |
| **SE** | 85.27 | 15.22 |
| **SME** | 89.90 | 14.02 |
| **Stability** | 0.90 | 0.09 |

*Note.* TIB = Time in Bed. SPT = Sleep Period Time. WASO = Wake After Sleep Onset. TST = Total Sleep Time. N1, N2, N3, and REM: Sleep stages duration. NREM = N1 +

N2 + N3. SOL = Sleep Onset Latency. Lat_N1, N2, N3, REM: latencies of sleep stages from the beginning of the record. %(N1, N2, N3, REM): Sleep stages duration expressed in percentages of TST. S.E. = Sleep Efficiency. SME: Sleep Maintenance Efficiency. Stability = diagonal value in the transition matrix.
